# IMAS resolves perturbation-sensitive regulatory architectures from matched tumour multiomics

**DOI:** 10.64898/2026.04.09.717444

**Authors:** Wu Deyang, Takashi Yamashiro, Toshihiro Inubushi

## Abstract

Tumour single-cell datasets contain weak, sparse and context-restricted regulatory signals that are difficult to distinguish from noise using expression measurements alone. Here we present IMAS, an integrative multiomic augmentation system that learns transferable regulatory structure from a pan-cancer foundation of matched single-cell RNA and chromatin-accessibility profiles and adapts it to data-limited target datasets. We refer to the resulting target-specific regulatory architectures as multi-layer target dependencies (MLTDs). MLTDs prioritize signals that remain supported across coordinated molecular, communication and perturbation-sensitive evidence, rather than by expression magnitude or any single prediction score. Rather than replacing the observed expression matrix, IMAS adds an interpretable regulatory-support layer for mechanism discovery.

Across independent tumour datasets, IMAS preserved matched cross-layer supervision, concentrated predictive support into compact target-aligned structures and improved recovery of RNA and transcription-factor states. In colorectal cancer, MLTDs resolved a SOD2-associated perturbation-sensitive architecture that was distinct from native expression, conventional co-expression and the inherited pan-cancer hierarchy. These dependencies promoted propagation of perturbation-associated information through RNA–TF–regulatory-element bridges and into receiver-TF-aware communication across malignant cell states. A LAMB1-centred analysis further showed that successive regulatory, communication and temporal constraints restored an expected extracellular-matrix programme that was weakly represented in the original matrix. In head and neck squamous cell carcinoma, SOX2-centred MLTDs resolved malignant-state-specific perturbation programmes. In renal cancer, MLTD-guided analysis identified tumour–vascular coupling associated with spatially localized endothelial-to-mesenchymal-transition-like states in Xenium data.

Together, IMAS reframes tumour multiomic augmentation as the recovery of compact, target-specific regulatory architectures rather than expression-matrix completion, providing an interpretable framework for prioritizing experimentally tractable mechanisms in heterogeneous tumour systems.

## 1. Introduction

Single-cell multiomic profiling enables transcriptional states and chromatin accessibility to be measured in the same cells, providing a direct basis for linking gene expression to regulatory-element usage and transcription-factor activity^1–3^. However, matched RNA–chromatin datasets remain much smaller and less diverse than single-cell RNA-sequencing atlases^4–6^. This limitation is particularly consequential in tumours, where regulatory signals may be weak, restricted to rare cellular states or confined to local microenvironmental interactions^7–9^. Such signals can be missed by expression-centred analyses, whereas unconstrained recovery risks converting technical variation into apparently coherent regulatory patterns^10,11^.

Existing computational approaches address individual aspects of this problem. Cross-modal models predict unmeasured molecular features^12,13^; regulatory-network methods infer TF–gene or enhancer–gene relationships^14–16^; ligand–receptor frameworks estimate potential intercellular communication^17,18^; and perturbation models predict transcriptional responses to genetic or pharmacological intervention^19,20^. These methods have substantially expanded single-cell analysis, but their outputs are generally evaluated separately. Consequently, a predicted regulatory element, TF response, communication event and perturbation effect are rarely organized within the same target-specific support structure. Moreover, increasing the amount of RNA-only training data does not necessarily preserve the matched cross-layer supervision required for regulatory traceability.

To address this gap, we developed IMAS, an Integrative Multiomic Augmentation System that learns transferable regulatory structure from pan-cancer matched RNA and chromatin-accessibility data and adapts it to data-limited target datasets. We refer to the resulting target-specific regulatory architectures as multi-layer target dependencies (MLTDs). An MLTD retains a candidate signal when it remains supported across complementary molecular or cellular layers, rather than because it is highly expressed, differentially expressed or assigned a high score by a single model. IMAS first learns transferable RNA–TF–regulatory-element relationships from matched multiomic profiles and then re-ranks and refines these relationships within the target cellular context.

IMAS therefore differs from conventional imputation and matrix-reconstruction workflows. It does not replace the observed expression matrix or attempt to generate a complete missing regulatory system. Instead, IMAS preserves the native target data and adds an interpretable MLTD layer that organizes candidate mechanisms according to matched multiomic support, target-specific adaptation, intercellular communication and perturbation-sensitive local structure. Augmentation thus refers to the addition of traceable regulatory support rather than substitution of the measured molecular state.

We used IMAS to determine what MLTDs capture and whether they support mechanism discovery across independent tumour systems. In colorectal cancer, MLTDs resolved a SOD2-associated perturbation-sensitive architecture that was distinct from native expression, conventional co-expression and the inherited pan-cancer hierarchy. These structures subsequently promoted propagation of perturbation-associated information through RNA–TF–regulatory-element bridges and into receiver-TF-aware communication across cellular states. A LAMB1-centred analysis further showed how successive regulatory, communication and temporal constraints restored a biologically expected extracellular-matrix architecture that was not prominent in the original matrix. In head and neck squamous cell carcinoma, SOX2-centred MLTDs resolved distinct malignant-state perturbation programmes. In renal cancer, MLTD-supported analysis identified tumour–vascular coupling associated with spatially localized EndMT-like endothelial states. Together, these analyses establish IMAS as a matched multiomic foundation and adaptation framework that converts sparse tumour signals into compact, target-specific and experimentally tractable regulatory hypotheses.

## 2. Materials and Methods

### 2.1 Study datasets, and preprocessing

#### Datasets and sample selection

Publicly available 10x Genomics multiomic tumour datasets were collected from open-access online resources (Table S1). Only datasets with matched multiomic measurements from the same cells were included. Tumour samples with sufficient data completeness for downstream analyses were retained, whereas datasets lacking paired measurements or containing incomplete matrices were excluded.

#### Preprocessing of RNA, TF, and regulatory features

RNA count matrices were generated from 10x Genomics matrix, feature and barcode files, retaining only features annotated as Gene Expression. Gene symbols were unified across samples to construct a global gene vocabulary, and each sample was aligned to this shared feature space. For cell*_i_*, RNA depth was defined as 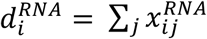, RNA depth was defined 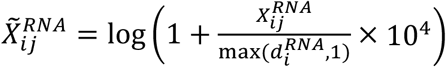. ATAC peak matrices were standardized by converting peaks to chromosome:start-end format and merging peaks across samples to define a global regulatory-element vocabulary. Sample-specific matrices were projected to the shared peak space, duplicate mappings were summed, and the resulting matrix was binarized as 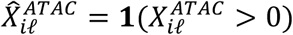. TF features were defined as the intersection between the global gene set and a curated human TF reference list^21^ 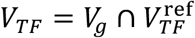. Gene coordinates, including chromosome, transcription start site and strand, were extracted from the gene annotation file, and a unified node index was constructed for TF, regulatory-element and gene nodes. Cells were matched across RNA and ATAC modalities using exact matches of normalized sample identifiers and shared cellular barcodes.

#### Construction of prior edges and spatial mapping

A multi-relation prior graph was constructed by integrating TF–RE, RE–gene and derived TF–gene edges. TF–RE edges were obtained by scanning peak sequences with FIMO from the MEME Suite using a curated motif collection in MEME format (motifs.meme) ^22,23^, and were then reweighted by a rank-evidence scheme 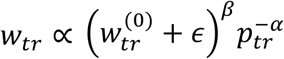, followed by normalization within each TF. Peak coordinates were unified in the hg38 reference space; when necessary, hg19 peak coordinates were lifted over to hg38 before motif scanning^24,25^. RE–gene edges were then defined by genomic proximity, where each regulatory element was linked to up to three nearest genes within 200 kb, with distance-decayed weights 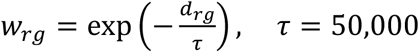. TF–gene prior edges were then assembled by composing TF–RE and RE–gene relations 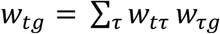, and the top 2,000 target genes were retained for each TF. All TF, regulatory-element and gene nodes were mapped into a unified graph space using the node index. Reverse edges were further generated by swapping source and destination nodes while preserving edge weights.

#### Global transcriptomic landscape

A shared RNA feature space was constructed across all datasets, followed by assembly of a global manifest indexing all cells. Feature genes were selected on the basis of cross-dataset frequency and expression variance in sampled cells. Expression values were normalized to 10,000 counts per cell and log-transformed 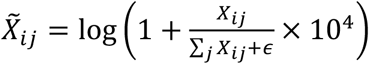. A stratified subset of cells was used to train Incremental PCA^26^ and UMAP^27^, and the fitted model was then applied to all cells to generate the global embedding.

### 2.2 Predictive modelling and transfer adaptation

#### Baseline predictive model construction

A baseline predictive model was constructed in the global pan-feature space after projecting all RNA and ATAC inputs, together with the six prior relation types, into a unified node index. Paired cells were assembled by exact matching of normalized sample identifiers and cellular barcodes. The model was trained in a leave-one-dataset-out (LODO) framework, in which one dataset was held out as the target domain and all remaining datasets were used for source training. For each paired cell, sparse RNA and RE inputs were truncated to fixed per-cell token budgets, where tokens correspond to retained non-zero projected features. In the baseline model, the RNA token budget was set to 2,048 features per cell and the RE token budget was set to 1,024 features per cell; in each case, the top-valued features were retained before encoding into the shared latent space. Model optimization combined cross-modal alignment with prior-graph supervision

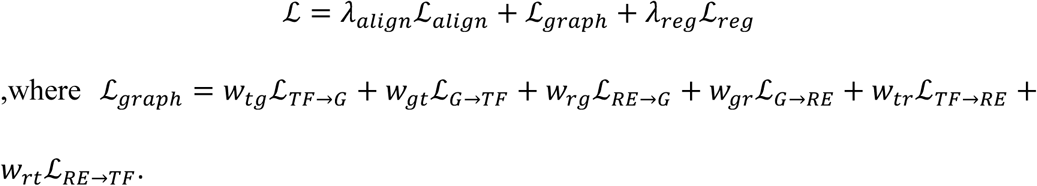

Training used AdamW with gradient clipping, and the resulting source-trained model was taken as the baseline before subsequent transfer adaptation^28,29^.

#### Transfer learning and target-domain adaptation

Target-domain adaptation was performed by freezing the matched multiomic foundation backbone and optimizing target-specific adaptation modules. RNA representations from the target dataset were encoded in the shared latent space and optimized through three transfer objectives: RNA-to-TF prediction, RNA-to-RE prediction and RNA-to-RNA reconstruction. Cell states in the target domain were further partitioned by spherical k-means, and group-specific adaptation was introduced through low-rank hyper-adapters and cluster-aware correction modules. To enhance target-specific regulatory signals, a selected-peak residual branch was added on top of the hash-based peak representation. The overall transfer objective was

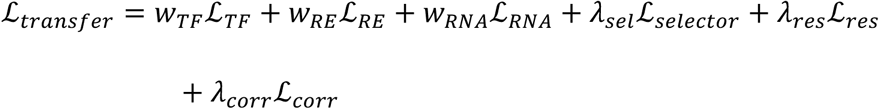

hyperparameters were fixed across experiments after preliminary tuning based on validation performance. The final settings were

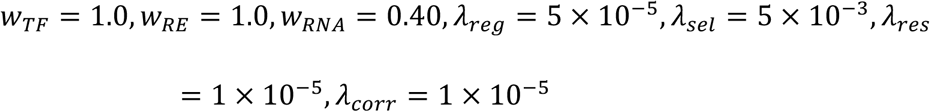

The TF and RE margin parameters were both set to 0.2, the adaptor rank was set to 32, and optimization used AdamW for 10,000 steps with a learning rate of 7 × 10^−4^ . Model selection was based on validation TF AUC, and the best checkpoint was retained for downstream evaluation in the target sample.

### 2.3 Interpretability analysis and predictive support validation

#### Predictive support identification for MLTD construction and counterfactual validation

Model predictive supports, representing the initial support elements for MLTD construction, were identified by extracting cell- and TF-specific explanatory subgraphs from the adapted model and comparing them with the baseline across RNA–TF, RNA– RE and TF–RE relations. Explanation patterns were further summarized by heatmaps and selection-contraction statistics to quantify concentration of positive mass and the effective number of selected nodes. For a given normalized importance vectorp, concentration was quantified as 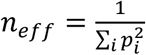. Faithfulness was evaluated by progressively deleting or retaining top-ranked predictive support genes and tracking the normalized target TF score. Stability was assessed by repeated perturbation of input tokens followed by overlap analysis of the top-ranked predictive supports. Counterfactual validation was performed by sequentially removing predictive support genes until prediction flipping or a marked score drop was observed, and the resulting trajectories were summarized across cases.

#### Trend consistency and manifold correspondence analysis

To assess whether transfer adaptation preserved biologically meaningful variation patterns, predicted and true profiles were compared at the cluster level for both RNA and TF features. For each cluster pair, a trend score was computed by combining cosine similarity, Pearson correlation and direction agreement between ordered feature profiles *S_trend_* = *ω*_1_*S*_cos_ + *ω*_2_*S_Pearson_* + *ω*_3_*S_dir_* . Trend, marker and structure scores were further integrated into a unified cluster correspondence score

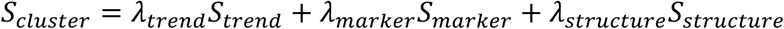

Cluster-level similarity and identity matrices were then constructed to quantify agreement between predicted and true cluster profiles. Cells with positive trend recovery were mapped back onto the original UMAP manifold to evaluate whether trend-consistent predictions remained aligned with the underlying cellular structure.

### 2.4 Stable regulatory network construction

#### Construction of stable RNA predictive network

These predictive RNA and TF networks were used as intracellular support layers for subsequent MLTD construction. To evaluate the stability of RNA predictive structure in the target domain, RNA-to-RNA retrieval performance was assessed using the adapted cell embeddings and gene embeddings in the unified latent space. For each cell, highly expressed genes were treated as positives and expressed but non-top-ranked genes as hard negatives, and gene relevance was scored by the dot product between the corrected cell embedding zi and gene embedding 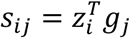. Cell-level retrieval performance was quantified by the area under the ROC curve (AUC), and mean ROC curves were aggregated across cells and across newly defined Leiden clusters obtained from the true RNA manifold^30^. Cluster-specific predictive stability was then summarized by comparing RNA-to-RNA retrieval performance across these reconstructed cell states.

#### Construction of stable TF predictive network

To evaluate the stability of TF predictive structure in the target domain, RNA-to-TF retrieval performance was assessed using adapted cell embeddings and TF embeddings in the shared latent space. For each cell, TF relevance was scored by the dot product between the corrected cell embedding zi and TF embedding tj 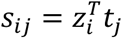. Cell-level prediction performance was quantified by the area under the ROC curve (AUC), with positive and negative TF labels defined from proxy TF activity profiles. Mean ROC curves were then aggregated across all cells and across newly defined Leiden clusters reconstructed from the true RNA manifold, thereby summarizing cluster-resolved TF predictive stability.

#### Assembly of the unified RNA–TF coupling network

A unified RNA–TF coupling network was assembled for each target sample using the adapted cell embeddings, gene embeddings and TF embeddings in the shared latent space. Candidate RNA–TF pairs were scored by integrating three sources of evidence: the Spearman association between RNA expression and predicted TF scores across cells, the latent gene–TF edge weight, and support from the TF–gene prior graph. For gene g and TF t, the network-aware coupling score was defined as

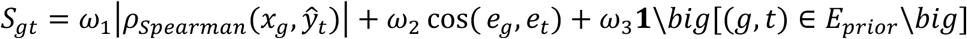

Scores were further rescaled to [0,1] and assembled into an RNA–TF score matrix, which was reordered to concentrate globally high-scoring interactions for visualization as the unified coupling heatmap.

### 2.5 Cell–cell communication modelling and communication-regulatory MLTD refinement

#### Communication-informed MLTD reinforcement using ligand – receptor information

Cluster-level ligand and receptor activities were derived from a combined gene-evidence term, obtained by z-scoring model-derived scores and cluster-mean log1p expression across clusters and averaging the two. The resulting evidence values were mapped to (0,1)by a sigmoid transform to define *E_s_*(*l*) *and E_r_*(*m*). Receiver-side TF response was z-scored across clusters and converted to a one-sided gating term *G_r_*(*m*) ∈ (0,1], which equalled 1 for non-negative response and decayed exponentially for negative response. The final communication score 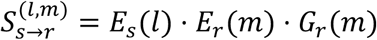 therefore also lay in (0,1) and was used for within-sample ranking of candidate interactions.

#### GNN–LTC modelling of communication pseudotime

A cluster-level communication graph was constructed from high-confidence cell– cell interactions, with nodes representing newly defined Leiden clusters and directed edges representing sender-to-receiver communication. Each node was characterized by sender strength, receiver strength, net flow, input–output ratio, receiver-side TF response and the centroid latent embedding of its member cells. Communication pseudotime was then learned using a GNN–LTC framework, in which graph-based neighborhood aggregation was combined with continuous-time latent rollout^31–33^. A weak teacher signal was first defined as

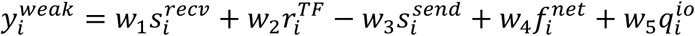

Where 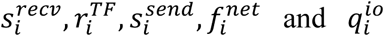 denote receiver strength, receiver TF response, sender strength, net flow and input–output ratio, respectively. The learned pseudotime was optimized by combining teacher fitting, rank consistency, graph smoothness, directed-edge consistency and feature reconstruction

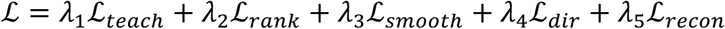

The resulting cluster pseudotime was further projected to individual cells by neighborhood smoothing in the latent space, enabling joint analysis of communication flow at the cluster, edge and cell levels.

#### Static projection of communication pseudotime and cluster communication states

To visualize the inferred communication program on a fixed cellular manifold, cluster-level communication pseudotime outputs were projected onto the single-sample reclustered UMAP without recomputing the embedding. Cluster-level metrics, including normalized pseudotime, uncertainty, LTC timescale, sender strength, receiver strength, net flow, input–output ratio and receiver-side TF response, were assigned to member cells according to their cluster identity. In parallel, cell-level projected pseudotime was directly overlaid onto the same manifold. This procedure enabled static visualization of communication states and pseudotime progression while preserving the original within-sample cellular geometry.

#### Dynamic projection of sender, receiver, and TF communication states

To visualize communication dynamics on the cellular manifold, sender strength, receiver strength and receiver-side TF response were projected onto a fixed single-sample UMAP using the inferred communication pseudotime. Cluster-level communication scores were first assigned to member cells and aligned along pseudotime, with sender, receiver and TF states shifted sequentially to reflect cascade ordering. For celli at pseudotime ti, the dynamic intensity of state k∈{sender,receiver,TF} was weighted by a Gaussian kernel 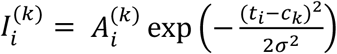, where Ai(k)denotes the normalized state score, ck the aligned state centre and σ the temporal bandwidth. The three state channels were then combined as an RGB cascade on the UMAP manifold, enabling dynamic visualization of sender-to-receiver-to-TF progression, together with cluster-level bubble summaries of the corresponding communication scores.

#### Identification of ligand–receptor temporal programs

Ligand–receptor temporal programs were identified by integrating post-pseudotime communication edges with receptor-specific TF context. For each sender– receiver cluster pair, the original ligand–receptor score was temporally reweighted according to the pseudotime shift between receiver and sender states 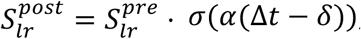, where Δt = *t_receiver_*-*t_sender_*, σ denotes the sigmoid function, and α and δ control temporal sharpness and delay. High-confidence ligand–receptor pairs were then ranked by temporally weighted scores and summarized in a TF-aware LIANA-style dotplot^34^, in which dot size reflected reinforced interaction strength and colour encoded the dominant receiver-side TF program. To visualize temporal reorganization at the network level, pre- and post-modelling cluster communication matrices were further aggregated and displayed as circle plots ordered by communication pseudotime, enabling identification of ligand–receptor interactions that were selectively retained, suppressed or temporally aligned after pseudotime refinement.

#### Pseudotime-resolved summarization and anchor-guided analysis of communication programs

Communication programs were further summarized after pseudotime modelling at both the cluster-feature and edge levels. Along cluster communication pseudotime, sender strength, receiver strength, receiver TF response, net flow and input–output ratio were quantified using pseudotime-resolved descriptors, including dynamic amplitude, coverage, maximal rate of change and midpoint timing. For a smoothed feature curve μ(t), these metrics were defined as

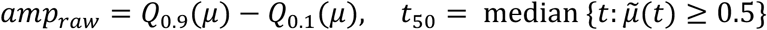

where ũ(t) denotes baseline-normalized feature intensity. Key sender–receiver edges were then ranked using edge weight, temporal shift and receiver-side TF response, and edges with similar temporal profiles were grouped relative to a seed edge by a composite similarity score combining profile correlation and timing proximity. In parallel, ligand–receptor programs were organized relative to a seed LR pair by binning interactions along sender pseudotime, smoothing pair-specific activity curves, and comparing their derivatives. For LR pairp, local agreement with the seed program was measured as 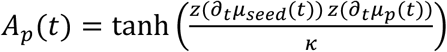, thereby enabling identification of LR programs that showed sustained agreement, divergence or opposite temporal behaviour relative to the anchor interaction.

### 2.6 LAMB1-centred network reinforcement and perturbation analysis

#### Stepwise reinforcement and in silico analysis of the LAMB1-centred MLTD

A LAMB1-centred local regulatory network was assembled in the shared latent space to characterize progressive signal reinforcement and functional dependency under in silico perturbation. Baseline node states were first defined from normalized expression and field strength, and effective edge weights were used to construct the initial interaction field. Network response to perturbation was then quantified by comparing post-perturbation and baseline node states and edge fluxes. For node*_i_*, the state change was defined as 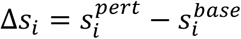 and for edge *u*→*v*, the flux change was defined as 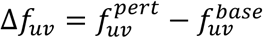 . These quantities were integrated to identify nodes and edges showing amplified, attenuated or redirected influence relative to the LAMB1-centred baseline field. The resulting stepwise network states were further visualized in three-dimensional space, in which node position encoded baseline topology and field intensity, while perturbation-induced changes were represented by node-state shifts and edge-flux deviations, thereby revealing a perturbation-aware local MLTD architecture around the target gene.

#### Combinatorial perturbation space design and perturbation engine

A combinatorial perturbation space was designed around the target-centred local network to evaluate the internal consistency of the perturbation model. Candidate perturbation nodes were selected from the reinforced local graph according to node type and normalized expression strength, with single-gene knockout used as the atomic perturbation unit. All pairwise combinations among candidate nodes were then enumerated to define the double-perturbation space. A random subset of gene pairs was retained as observed combinations, whereas the remaining pairs were held out as unseen perturbations for internal evaluation. Formally, for a candidate set V={*g_1_*,…,*g_n_*}, the combinatorial space was defined as *P*_2_ = {(*g_i_*, *g*_j_) ∣ 1 ≤ *i* < *j* ≤ *n*} . This yielded three perturbation categories: diagonal single-gene perturbations, observed double perturbations, and predicted unseen double perturbations. The local perturbation engine was then used to infer the held-out combinatorial space from the observed subset, thereby establishing the setup for internal combinatorial perturbation assessment.

#### Visualization of single-gene perturbation effects on the UMAP manifold

Single-gene perturbation effects were visualized on a fixed UMAP manifold constructed from the true RNA expression space. Baseline cell embeddings were first obtained by normalization, log transformation, principal component analysis and UMAP projection. For a target gene, a virtual knockout program was then defined from local gene–gene correlations, and perturbed expression states were generated by propagating the knockout effect to correlated genes in a cell-dependent manner. The perturbed states were projected back to the original UMAP through the baseline PCA space and k-nearest-neighbour interpolation. For cell*_i_*, the manifold displacement was defined as 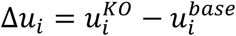. Cluster-weighted transcriptional responses were further aggregated into a grid-based vector field on the UMAP manifold to visualize the direction and magnitude of perturbation-induced shifts.

### 2.7 External perturbation benchmarking and SOX2-centred analysis

External perturbation benchmarking was performed using perturbation-associated reference genes as directional response labels. IMAS-predicted response directions were compared with CPA, scGPT and GEARS, and same-trend recovery was defined as the fraction of reference-supported genes for which the predicted direction matched the observed perturbation-associated direction. Local MLTD connectivity and component-ablation analyses were used to assess whether response recovery depended on the integrated IMAS-derived MLTD. For independent target-centred analysis, an HNSCC single-cell RNA-seq dataset was processed by standard quality control, normalization, dimensionality reduction, clustering and marker-based annotation. SOX2 expression was quantified across sample groups and projected onto the cellular manifold. A SOX2-centred MLTD was constructed by integrating target-context expression support, prior/TF regulatory support and gene-node coupling evidence. Virtual SOX2 knockout was performed by propagating an initial SOX2 perturbation signal through this SOX2-centred MLTD. Cell-level response scores were summarized across annotated cell types, projected onto the fixed UMAP manifold and used to define perturbation phases. Phase-associated ligand–receptor communication and malignant-cell response states were then analysed to identify SOX2-associated response programs.

### 2.8 Patient-derived renal tumour single-cell cohort and MLTD-guided matrix-preserving augmentation

Fresh renal tumour tissues were collected from eight patients undergoing surgical resection at Gifu University, comprising four clear-cell renal cell carcinomas (ccRCCs), three chromophobe renal cell carcinomas and one papillary renal cell carcinoma. Histological diagnoses were confirmed by pathological examination. The study was approved by the institutional ethics committee (approval number:), and written informed consent was obtained from all participants.

Fresh tissues were mechanically and enzymatically dissociated into single-cell suspensions. Single-cell RNA-sequencing libraries were prepared using the 10x Genomics Chromium platform. Reads were aligned to the GRCh38 reference genome and quantified using Cell Ranger [v10.1.0]. Low-quality cells, putative doublets and cells with high mitochondrial transcript fractions were removed using sample-specific quality-control thresholds. The eight samples were normalized and integrated in a shared gene-expression space. Principal-component analysis, neighbourhood-graph construction, Leiden clustering and UMAP were used to resolve the cellular landscape. Cell identities were assigned from cluster-enriched genes and canonical lineage markers, resolving tumour epithelial, tubular epithelial, endothelial, pericyte, lymphoid, myeloid and stromal populations. Endothelial-cell and pericyte numbers were summarized separately for each sample.

In parallel with the native expression UMAP, cells were represented using their IMAS-derived MLTD support profiles. These profiles summarized target-adapted regulatory and network support associated with each cell and were projected into a separate low-dimensional representation, termed the MLTD-echo UMAP. This representation was used to assess whether sample and lineage structures were retained in the dependency-supported space and did not alter or replace the original expression matrix. The IMAS-mapped profiles from all eight renal tumour samples were compared with a healthy-kidney single-cell reference to define a cohort-level tumour differential landscape^35^. Among the eight tumours, Sample 2 contained the largest endothelial-cell compartment and was therefore selected as the target ccRCC sample for subsequent MLTD-guided tumour–vascular analysis. The remaining samples were retained for cohort-level atlas construction and differential-state comparison but were not pooled into the target-specific communication and functional-coupling analyses. Within the Sample 2 target context, an ordinary DEG set relative to the healthy-kidney reference was retained as the native differential backbone.

IMAS network-differential scores integrated expression differences with graph-position shifts, propagation-field changes and MLTD structural support. Genes outside the ordinary DEG set were retained as MLTD-supported incremental nodes when they satisfied the predefined network-support criteria. The augmented differential set was defined as the union of the unchanged DEG core and these traceable incremental nodes. Functional enrichment^36–39^, ligand–receptor communication and tri-constraint coupling analyses were then performed to connect tumour expression states, MLTD-supported differential nodes and communication constraints. Ablation analyses removed expression, differential or communication constraints from the full model. IMAS was further compared with MAGIC^40^ and ALRA^41^ workflows by evaluating whether downstream DEG sets were generated after matrix replacement or by preserving the raw DEG core with a traceable MLTD-supported increment.

### 2.9 ccRCC Xenium spatial validation

An independent public Xenium dataset, “Xenium In Situ Gene and Protein Expression data for FFPE Human Renal Cell Carcinoma” from 10x Genomics, was used to evaluate spatial support for the IMAS-nominated tumour–vascular program. The dataset was generated from FFPE human renal cell carcinoma tissue using the Xenium In Situ Gene and Protein Expression with Cell Segmentation Staining workflow and was analysed with Xenium Onboard Analysis v4.0.0^42^. Cell-level transcript counts and spatial coordinates were processed by quality control, normalization and marker-based annotation. Tumour epithelial, endothelial, immune, stromal and proliferative compartments were identified using canonical markers. Endothelial structures were marked by PECAM1 and ERG, EndMT-like features by ACTA2 and VIM, vascular-abnormality programs by ANGPT2, ADGRL4 and VWF, and proliferative epithelial neighbourhoods by MKI67, PCNA and KRT8. Spatial maps were generated by overlaying marker or module scores onto cell coordinates. EndMT-like vascular regions were defined as endothelial-associated neighbourhoods with elevated ACTA2/VIM signal. Endothelial cells were further stratified into ACTA2/VIM-high and ACTA2/VIM-low groups for differential expression and pathway enrichment analysis. Spatial neighbourhood analysis was used to assess whether endothelial remodelling regions were located near tumour epithelial regions with RTK-associated and proliferative programs. These analyses were interpreted as spatial support for the IMAS-nominated tumour–vascular MLTD.

## 3. Results

### 3.1 Matched tumour multiomics establishes transferable and interpretable regulatory dependencies

Matched single-cell RNA and chromatin-accessibility measurements directly connect transcriptional states with their underlying regulatory potential, but such datasets remain substantially less abundant than scRNA-seq atlases^43,44^. This imbalance is particularly consequential in tumours, where regulatory relationships are frequently restricted to specific lineages, malignant states or microenvironmental contexts and may therefore be weakly represented in expression data alone^45–47^. We reasoned that the principal value of matched tumour multiomics is not simply the availability of an additional molecular modality, but the paired cross-layer supervision required to learn regulatory relationships that remain traceable across RNA, transcription-factor and regulatory-element spaces.

We therefore assembled a pan-cancer foundation corpus composed exclusively of matched single-cell RNA–ATAC tumour datasets spanning diverse cancer types and experimental cohorts, and trained IMAS in a unified gene, TF and regulatory-element feature space (Fig. 1). The shared foundation model jointly optimized RNA–ATAC alignment, cross-modal reconstruction and graph-aware link prediction, thereby learning transferable support for RNA–TF, RNA–RE and RNA–RNA relationships. Rather than treating RNA and chromatin accessibility as independently observed feature collections, IMAS used their cell-matched correspondence to constrain a shared regulatory representation. This design was intended to preserve the molecular traceability required for downstream mechanism inference, rather than merely maximize latent-space alignment or missing-modality prediction.

**Fig. 1.**
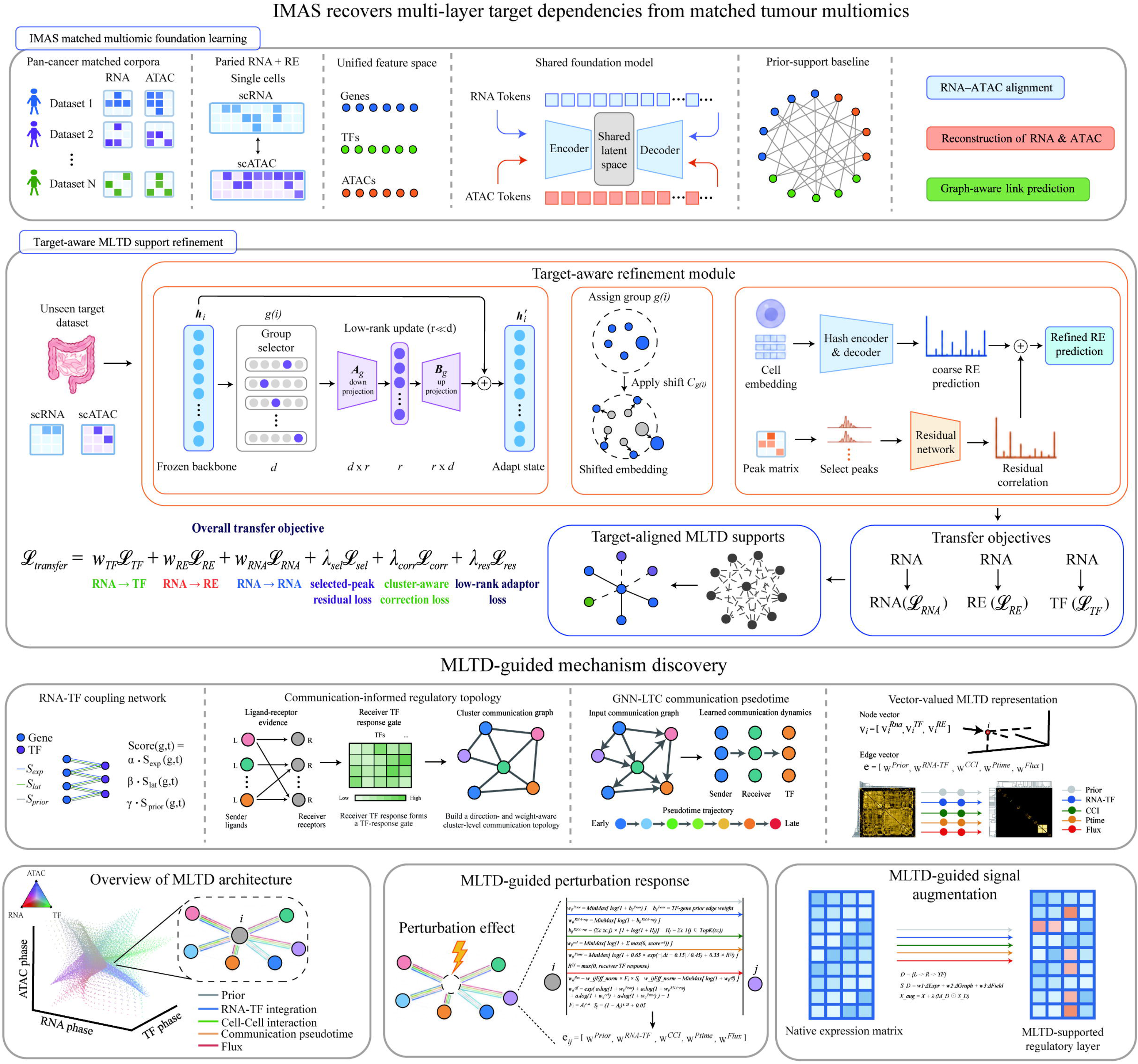
IMAS learns transferable multi-layer dependencies and refines them into target-aligned MLTD supports. Pan-cancer matched RNA and ATAC profiles are mapped into a unified gene, TF and regulatory-element feature space and used to train a shared foundation model through RNA–ATAC alignment, cross-modal reconstruction and graph-aware link prediction. For transfer to an unseen target dataset, the frozen foundation backbone is adapted using group-selective low-rank updates, cluster-aware embedding correction and selected-peak residual refinement. The transfer objective jointly constrains RNA–TF, RNA–RE and RNA–RNA prediction while preserving target-cluster structure and regulatory-element residuals. The resulting target-aligned MLTD supports are integrated with ligand–receptor evidence, receiver-TF responses, communication topology, GNN–LTC communication pseudotime and perturbation field/flux information to construct vector-valued node and edge representations for perturbation-response inference and augmentation of target regulatory layers.

The intended application of this foundation was the analysis of tumour datasets for which only scRNA-seq measurements are available. To transfer matched multiomic regulatory structure into such data-limited target systems, IMAS retained the shared foundation backbone and introduced group-selective low-rank adaptation, cluster-aware embedding correction and residual regulatory-element refinement (Fig. 1). These target-specific modules restricted adaptation to a compact parameter subspace while jointly constraining RNA–TF, RNA–RE and RNA–RNA prediction and preserving selected-peak residuals and target-cluster organization. The resulting target-aligned supports were subsequently integrated with ligand–receptor evidence, receiver-side TF responses, communication topology and GNN–LTC pseudotime to construct vector-valued multi-layer target dependencies (MLTDs) for perturbation inference and matrix-preserving regulatory augmentation.

The foundation corpus showed broad, although uneven, representation across tumour types, with medulloblastoma, prostate cancer and glioma contributing the largest numbers of matched cells (Extended Data Fig. 1a). The ten largest cohorts together accounted for 67.7% of cells, indicating substantial contribution from additional tumour and experimental contexts rather than dominance by a single disease group (Extended Data Fig. 1b). Joint embedding retained marked cohort, lineage and tumour-state diversity across the pan-cancer atlas (Extended Data Fig. 1c), providing a heterogeneous basis from which transferable regulatory relationships could be learned. We next tested whether matched multiomic supervision contributed a form of regulatory organization that could not be recovered simply by increasing the amount of RNA-only training data. Across colorectal cancer, glioma and osteosarcoma, foundations trained exclusively on matched multiomic cells produced stronger target dependency organization than foundations trained with mixed multiomic and RNA-only data or with RNA-only data alone (Extended Data Fig. 1d). This advantage was consistently observed across four complementary properties: overall dependency support, regulatory-element traceability, preservation of the dependency core and anchoring of perturbation-associated responses (Extended Data Fig. 1e). These differences therefore extended beyond predictive accuracy and affected whether inferred signals remained organized into compact, cross-layer and biologically traceable structures.

To directly assess the dependence of these properties on paired supervision, we progressively diluted matched multiomic information while retaining increasing amounts of RNA-only information. Reduction of the matched fraction caused a coordinated decline in dependency support, regulatory-element traceability, dependency-core preservation and response anchoring across tumour types (Extended Data Fig. 1f). Thus, larger quantities of transcriptomic data did not substitute for cell-matched cross-layer supervision. Instead, the paired RNA–chromatin relationship was the defining source of the regulatory structure required to distinguish weak but consistently supported dependencies from expression-associated noise.

Together, these results establish a central design principle of IMAS: for mechanism-oriented tumour foundation learning, the critical resource is not transcriptomic scale alone, but matched multiomic supervision that preserves cross-layer regulatory traceability. IMAS transfers this structure into RNA-only target datasets and refines it according to the local tumour context, producing target-aligned MLTDs for subsequent perturbation, communication and mechanism-discovery analyses.

### 3.2 MLTD resolves perturbation-sensitive dependency structures

Cellular responses to perturbation are shaped by context-dependent regulatory relationships that are not fully captured by expression abundance or pairwise correlation alone^48–50^. We therefore asked whether MLTDs could resolve the dependency architecture through which a defined perturbation is organized in a target cellular system. We used SOD2 as a representative perturbation seed because its established role in mitochondrial redox regulation provided an interpretable context for testing whether MLTD could recover biologically coherent perturbation-associated structure beyond expression abundance. In colorectal cancer, SOD2-centred MLTD identified a heterogeneous response architecture across malignant epithelial states and localized propagation domains on the cellular manifold (Fig. 2a–c). This organization was stable across potential thresholds, and the integrated propagation area under the curve exceeded matched randomized structures (P = 0.03792; Fig. 2d,e), indicating that the inferred field was more spatially organized than expected from matched null propagation. Native SOD2 expression differed only modestly between MLTD-high and MLTD-low cells (standardized mean difference, SMD = 0.14), whereas the inferred directional response was strongly separated (SMD = 1.86; Fig. 2f,g). The resulting states were enriched for oxidative-stress, respiratory-chain, ROS-detoxification, NRF2-response and mitochondrial-quality-control programmes (Fig. 2h), consistent with established functions of SOD2 in mitochondrial redox regulation^51,52^.

**Fig. 2.**
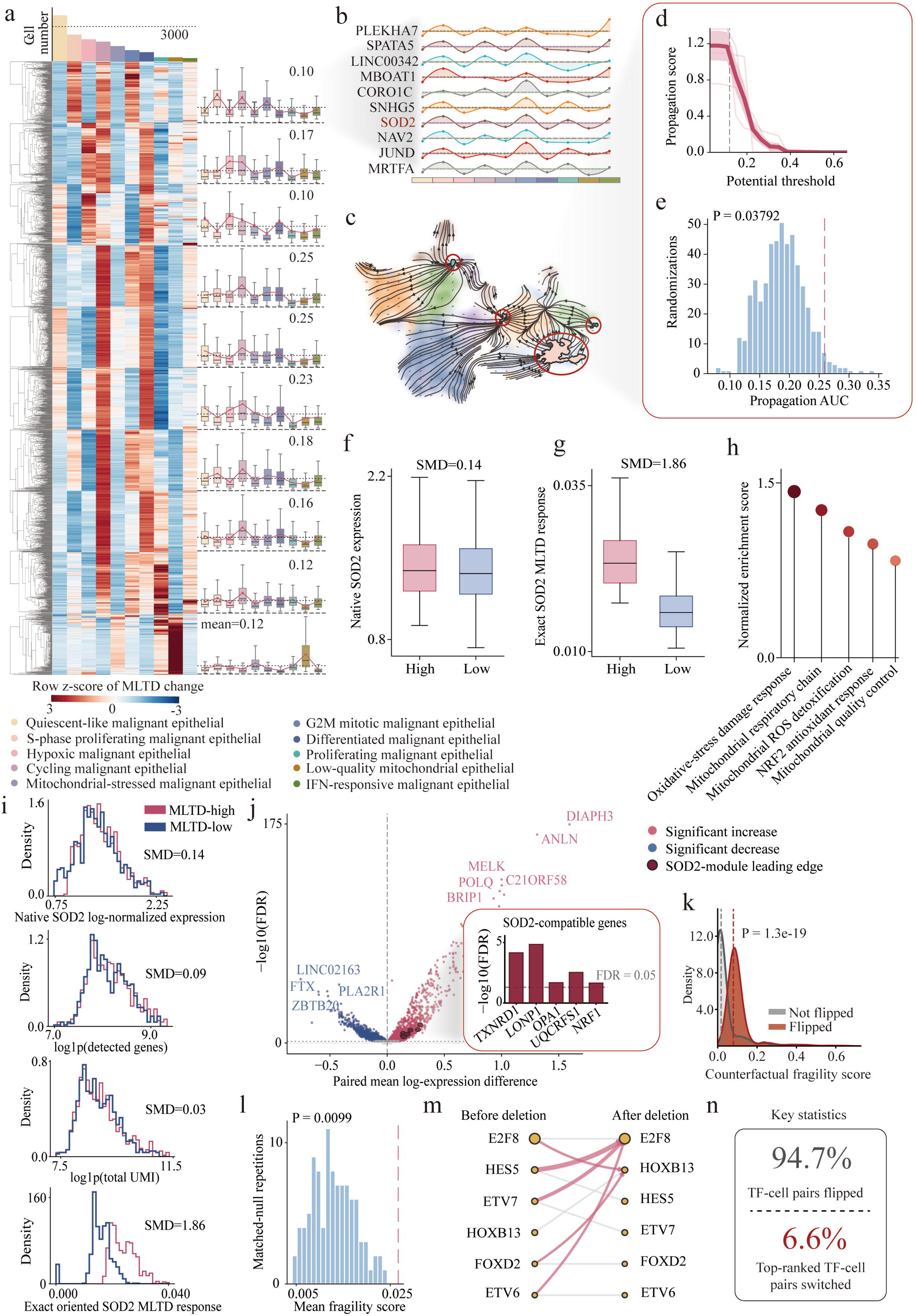
MLTD resolves perturbation-sensitive dependency structures in colorectal cancer. **a,** Heat map showing cell-level SOD2-centred MLTD response changes across 3,000 malignant epithelial cells ordered by cell state. Rows represent response-associated features and columns represent cells; values are row-wise z scores. Right, state-stratified distributions of MLTD changes. Colours denote malignant epithelial cell states. **b,** Cluster-resolved profiles of representative genes associated with the inferred SOD2 dependency structure, including SOD2. Curves show normalized values across malignant epithelial states; the colour strip indicates cell-state identity. **c,** Projection of the inferred SOD2 dependency-propagation field onto the malignant-cell manifold. Streamlines indicate local propagation directions; red circles highlight representative convergence or transition regions. **d,** Propagation score as a function of the potential threshold. The solid line shows the observed mean and lighter lines indicate repeated estimates. **e,** Null distribution of propagation area under the curve (AUC) obtained from matched randomizations. The dashed line indicates the observed value; the empirical P value is shown. **f,** Native SOD2 expression in cells with high or low MLTD dependency scores. **g,** Exact oriented SOD2 MLTD response in the same groups. SMD, standardized mean difference. **h,** Enrichment scores of mitochondrial and oxidative-stress programmes associated with the inferred SOD2 dependency structure. **i,** Density distributions of native SOD2 expression, detected gene number, total UMI count and exact oriented SOD2 response in MLTD-high and MLTD-low cells. **j,** Paired mean log-expression differences between MLTD-high and MLTD-low cells. Red and blue points indicate significantly increased and decreased genes, respectively. Inset, false-discovery rates for representative SOD2-compatible genes. **k,** Counterfactual fragility-score distributions for TF–cell pairs whose predicted direction was or was not flipped after rationale deletion. The P value was calculated using a two-sided comparison between groups. **l,** Null distribution of the mean fragility score from matched repetitions. The dashed line indicates the observed value; the empirical P value is shown. **m,** Changes in representative TF–cell rankings before and after rationale deletion. Line width reflects the magnitude of reassignment. **n,** Summary statistics showing the proportions of TF–cell pairs with directional flipping and top-ranked TF switching.

This separation was not explained by baseline expression or technical covariates. MLTD-high and MLTD-low cells remained closely matched in native SOD2 expression, detected gene number and total transcript abundance, but differed markedly in the inferred SOD2 response (Fig. 2i). Genes associated with the recovered structure included TXNRD1, LONP1, OPA1, UQCRFS1 and NRF1 (Fig. 2j), which collectively link antioxidant defence, mitochondrial proteostasis and respiratory-chain regulation^53–56^. Counterfactual deletion of the selected rationale preferentially destabilized TF–cell relationships that showed directional flipping (P = 1.3 × 10^−19^) and produced greater fragility than matched null deletions (P = 0.0099; Fig. 2k,l). Overall, 94.7% of TF–cell pairs showed directional flipping, whereas 6.6% switched their top-ranked TF, indicating broad reorganization of relative regulatory support without universal replacement of the dominant regulator (Fig. 2m,n).

The identified structure generalized across colorectal cancer, glioma and osteosarcoma. Retaining MLTD-prioritized relations preserved model support and cellular-state coverage, whereas deleting them caused larger losses than baseline or weight- and degree-matched null selections (Extended Data Figs. 2 and 3). These effects extended across RNA–TF, RNA–RE and TF–RE layers, indicating that the recovered response depended on coordinated cross-layer relations rather than a single interaction class.

**Fig. 3.**
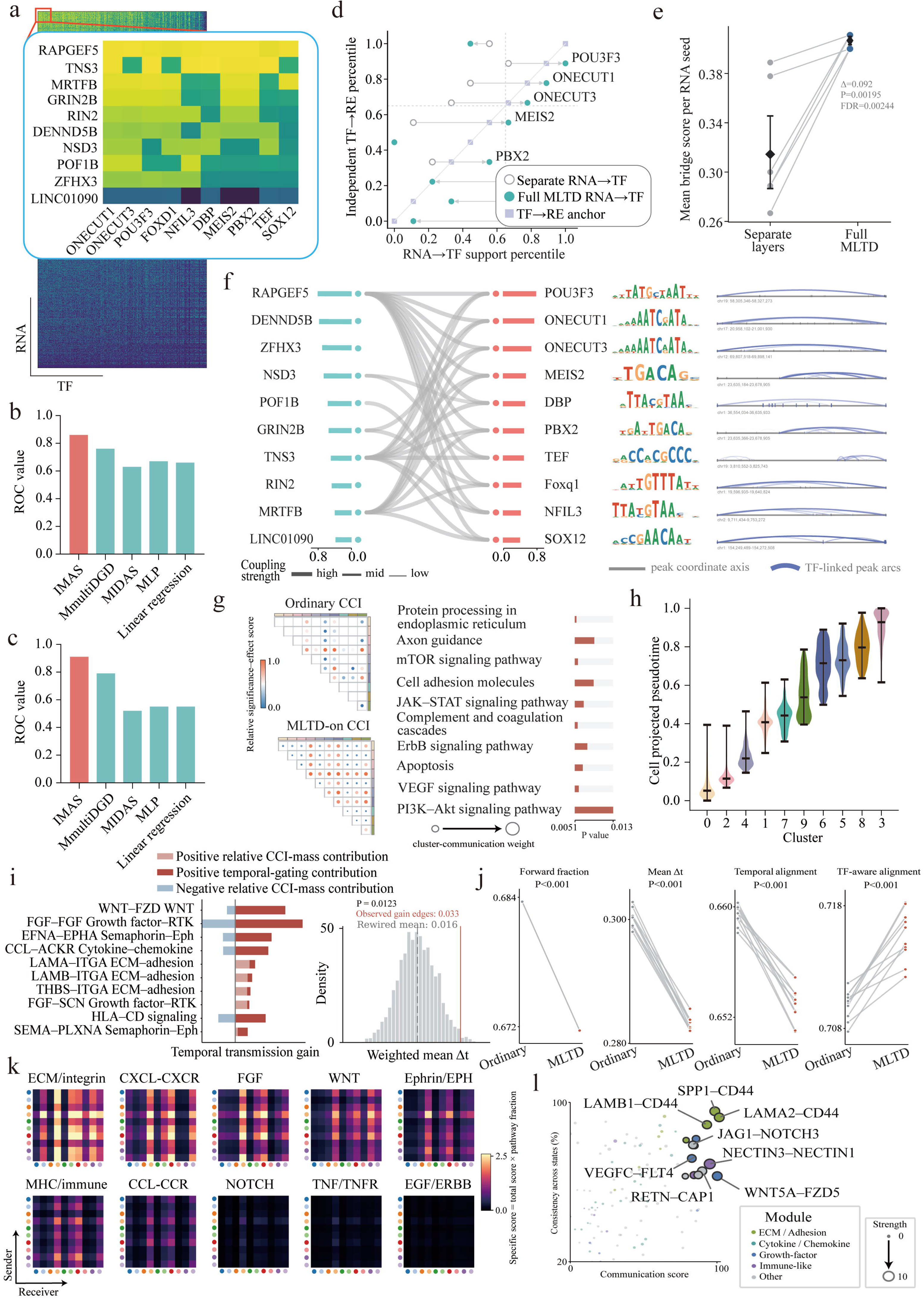
MLTD promotes perturbation propagation across molecular layers and cell states. **a,** Matrix of RNA-seed–TF bridge scores inferred by MLTD. The enlarged region shows representative RNA seeds and TF programmes with high bridge support. **b,c,** Receiver-operating-characteristic performance for RNA–TF inference across the indicated methods in two evaluation settings. **d,** Within-layer support percentiles for representative TF programmes obtained using separately inferred RNA–TF relations, full MLTD RNA–TF integration and TF–RE anchors. Lines connect estimates for the same TF. **e,** Paired comparison of mean RNA-seed bridge scores obtained from separate molecular layers and full MLTD. Points represent matched RNA seeds; the central point and error bars show the mean and uncertainty. The reported effect size, P value and false-discovery rate are shown. **f,**Representative RNA–TF–RE propagation paths. Left, RNA seeds; centre, TF programmes and corresponding DNA-binding motifs; right, genomic peak coordinates and TF-linked regulatory-element arcs. Line width indicates coupling strength. **g,** Cluster-level communication under ordinary CCI and MLTD-on CCI. Bubble size denotes cluster–communication weight and colour denotes relative significance–effect score. Right, the ten uniquely MLTD-gained pathways and their associated P values. **h,** Cell-projected communication pseudotime inferred from MLTD-informed sender–receiver communication and receiver-side TF response across the indicated clusters. Violins show cell-level distributions; central symbols and vertical bars show summary estimates and ranges. **i,** Pathway-level temporal-transmission gains after MLTD integration. Horizontal bars show positive temporal-gating contributions and positive or negative relative CCI-mass contributions. Right, null distribution of weighted mean receiver–sender time differences; dashed lines indicate the observed and rewired values. **j,** Paired comparisons of forward-edge fraction, weighted mean receiver–sender time difference, temporal alignment and TF-aware temporal alignment between ordinary and MLTD-informed communication. Lines connect matched model runs; P values are shown above each comparison. **k,** Ligand–receptor activity across receiver clusters ordered by fixed MLTD-off communication pseudotime. Heat maps are grouped by signalling module; colour denotes normalized activity. **l,** Communication score and consistency across cell states for representative ligand– receptor pairs. Bubble size denotes mean interaction strength and colour indicates signalling module. Labelled interactions are representative pairs from MLTD-gained pathways.

Finally, the SOD2-associated structure was not simply inherited from the pan-cancer foundation. MLTD dependency, but not native SOD2 expression or conventional co-expression, significantly recovered predefined mitochondrial programmes (P = 0.002, 0.115 and 0.234, respectively; Extended Data Fig. 4a,b). Foundation and adapted scores were substantially reordered across clusters and cells, with a global Spearman correlation of −0.42 and limited overlap among top-ranked cells (Jaccard index = 0.03; Extended Data Fig. 4c–f). Foundation adjustment introduced additional leading-edge genes and strengthened separation between MLTD-high and MLTD-low states (Extended Data Fig. 4g–i). Thus, MLTD resolves a compact, model-inferred perturbation-sensitive and target-specific dependency structure that organizes perturbation-associated information across molecular layers. Having established this dependency architecture, we next asked whether it could support propagation of perturbation-associated information across molecular layers.

### 3.3 MLTD promotes perturbation propagation across molecular layers and cell states

Perturbation responses are coordinated across transcriptional, regulatory and intercellular layers, yet these components are commonly modelled in isolation^57,58^. We therefore asked whether the perturbation-sensitive structures identified by MLTD could first recover target-cell RNA and TF states and then propagate perturbation-associated information across regulatory and intercellular layers. Across colorectal cancer, glioma and osteosarcoma, target adaptation improved RNA and TF profile concordance and preserved cellular-state geometry relative to the baseline, whereas removal of the low-rank adaptor or cluster correction reduced recovery (Extended Data Fig. 5a–d).

We next examined whether the recovered states were connected through coherent regulatory paths. MLTD improved RNA–TF prediction relative to alternative approaches and increased bridge support compared with independently inferred RNA– TF and TF–RE relations (Fig. 3a–e). These gains linked RNA seeds to TF programmes and motif-supported regulatory elements, forming coordinated RNA–TF–RE paths rather than isolated pairwise associations (Fig. 3f). The observed bridges exceeded TF– target rewiring controls and were concentrated within specific RNA-seed–TF-program combinations (Extended Data Fig. 5e–g), consistent with regulatory effects propagating through coupled molecular relationships^59–61^.

We then tested whether these cross-layer gains extended to communication between cell states. Relative to MLTD-off communication, MLTD-on CCI increased selected cluster-pair interaction and revealed pathway-specific gains involving PI3K– Akt, axon guidance, cell adhesion, JAK–STAT and VEGF signalling (Fig. 3g). These interactions connected ECM, chemokine, growth-factor, WNT, Ephrin and immune-response programmes across malignant states (Extended Data Fig. 5h,j). Because ligand–receptor activity alone does not specify the downstream regulatory response of the receiving cell^62,63^, we incorporated receiver TF activity into GNN–LTC modelling. GNN–LTC organized the MLTD-informed communication network along a communication pseudotime, reflecting sender-receiver flow and receiver-side TF response rather than a conventional transcriptomic trajectory (Fig. 3h). MLTD-on broadened the set of temporally supported interactions while largely preserving the underlying cluster order (Extended Data Fig. 5i). Pathway-level gains reflected both redistribution of CCI support and temporal gating, and MLTD selectively increased TF-aware temporal alignment despite limited improvement in global forward-direction measures (Fig. 3i,j). These effects exceeded graph-shuffled and receiver-TF-permuted controls across tumour types (Extended Data Fig. 6). Stage-wise analysis further showed that TF-target integration and CCI produced the principal gains in communication accuracy and TF-response recovery, whereas LTC organized these signals across cell states (Extended Data Fig. 7).

Together, these results show that MLTDs propagate perturbation-associated information from target-adapted RNA–TF–RE bridges into receiver-TF-aware communication between malignant cell states. Rather than acting as a generic domain-correction procedure, target adaptation re-ranks transferable regulatory support according to the target cellular context and enables cross-layer propagation of perturbation-sensitive regulatory information. We next asked whether such cross-layer propagation could progressively restore a dataset-specific local perturbation space around a representative communication-associated target.

### 3.4 MLTD progressively restores a dataset-specific perturbation space

Transferable priors may contain biologically relevant relationships that remain weak or undetectable in a target expression matrix because of sparsity, transcriptional heterogeneity and indirect regulation^64–66^. We therefore asked whether successive MLTD constraints could restore such latent relationships while adapting them to the target dataset. Using LAMB1 as a representative perturbation seed, representation, TF– RNA, communication, temporal and field/flux constraints progressively condensed a diffuse dependency field into modular local architectures (Fig. 4a–e). Across this progression, the top-response hierarchy was repeatedly reorganized: regulators supported by the prior were selectively retained or displaced, whereas factors including SCARB2, IL20RA, FAM47E, ERAP2, TAF15, MTF1L and LINC01011 gained support after target-derived constraints were introduced.

**Fig. 4.**
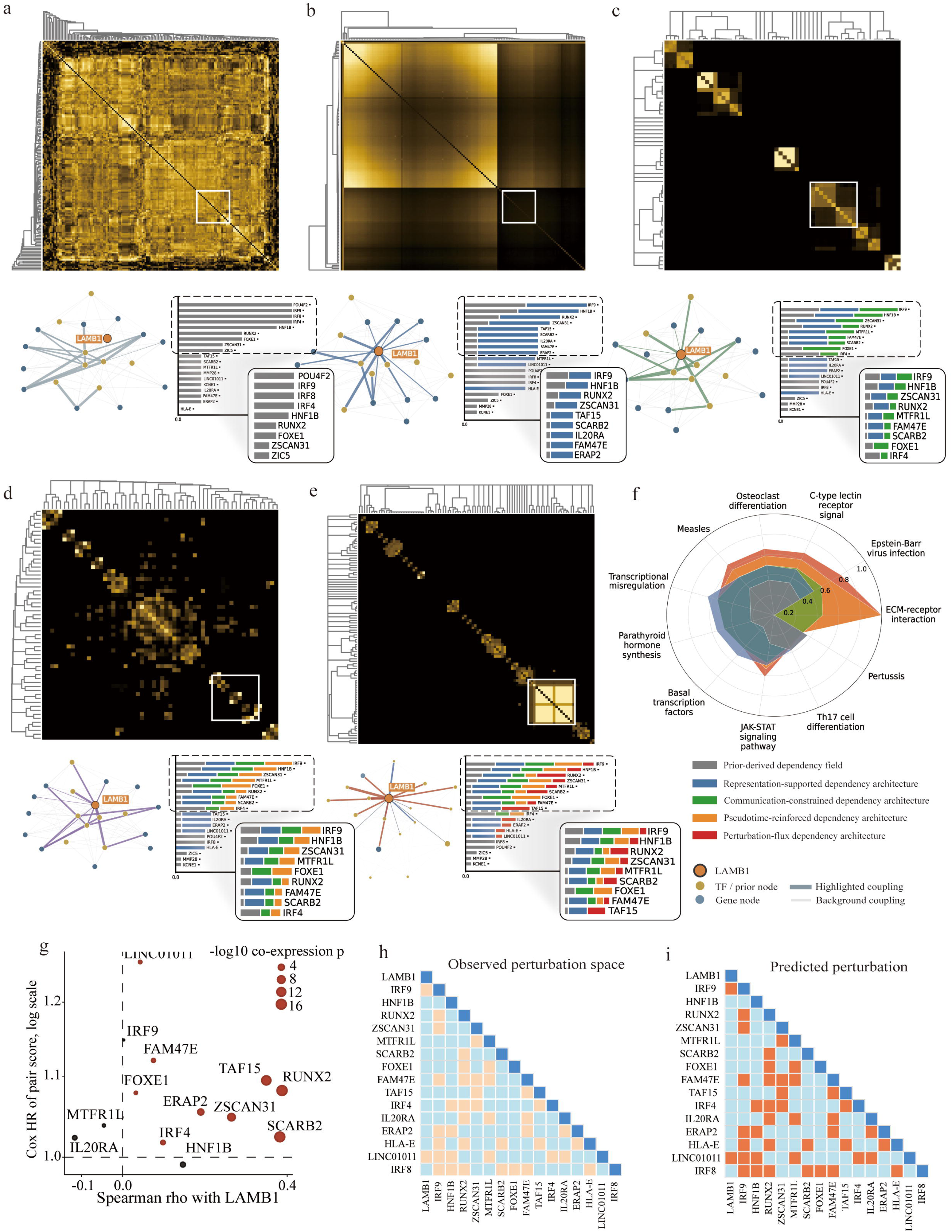
MLTD progressively restores a dataset-specific LAMB1 perturbation space. **a–e**, Progressive refinement of the LAMB1-centred dependency field after sequential incorporation of prior-derived support **(a)**, representation-supported structure **(b)**, communication constraints **(c)**, temporal information **(d)** and perturbation field/flux information **(e)**. Top, hierarchically clustered dependency matrices; white boxes mark representative local modules. Bottom, corresponding LAMB1-centred subnetworks and ranked response regulators. Node colour denotes node class, edge colour denotes the newly incorporated source of support and edge width indicates coupling strength. Insets show the highest-ranked predicted responders at each stage and illustrate progressive reordering of the response hierarchy. **f,** KEGG pathway support across the five dependency architectures. Radial profiles show normalized pathway recovery for the prior-derived, representation-supported, communication-constrained, pseudotime-reinforced and perturbation-flux architectures. The progressive recovery of ECM– receptor interaction highlights restoration of the expected LAMB1-associated extracellular-matrix programme. **g,** Relationship between co-expression with LAMB1 and pairwise dependency support for candidate responders. The x axis shows Spearman correlation with LAMB1, the y axis shows the Cox hazard ratio of the pair score on a logarithmic scale and point size denotes co-expression significance. Dashed lines mark zero correlation and a hazard ratio of 1. **i,** Internally observed pairwise perturbation-response structure among LAMB1 and the indicated regulators used for combinatorial perturbation assessment. **j,** Corresponding MLTD-predicted perturbation-response structure for inferred perturbation combinations. In i,j, colours indicate the direction and magnitude of pairwise response values.

A defining feature of this progression was the recovery of the expected relationship between LAMB1 and extracellular-matrix regulation. Laminins are core basement-membrane components that mediate cell–matrix adhesion and signalling through integrin and non-integrin receptors^67–69^. Nevertheless, ECM–receptor interaction was not prominently resolved in the initial prior-derived field. Its support increased progressively as representation, communication, temporal and perturbation-flux constraints were introduced, ultimately becoming the dominant pathway in the final architecture (Fig. 4f). This recovery occurred alongside repeated re-ranking of the highest-response regulators, indicating that MLTD did not simply amplify a predefined ECM prior but reorganized the target-supported routes through which the LAMB1-associated programme was represented.

This recovery did not reflect uniform amplification of all prior relationships. Although recurrent regulators such as IRF9, HNF1B, RUNX2, ZSCAN31, MTF1L and FOXE1 remained detectable across layers, their relative support was progressively redistributed (Extended Data Fig. 8a,b). The final field retained a similar fraction of nodes but concentrated a greater proportion of support within its strongest edges and reduced the effective support ratio (Extended Data Fig. 8c), indicating selective recovery rather than indiscriminate network densification.

The final dependency architecture was also not reducible to expression correlation. Several highly ranked responders showed little or no correlation with LAMB1, whereas others combined positive co-expression with high pairwise support (Fig. 4g). Because dropout and cellular heterogeneity can obscure regulatory relationships in single-cell expression data, and co-expression cannot distinguish direct from indirect dependencies^70,71^, we compared the inferred field with weight-permuted and degree-preserving rewired controls. These perturbations disrupted target-local strength, centralization and direct-to-indirect support ratios, placing the observed MLTD structure outside matched network configurations (Extended Data Fig. 8d).

Finally, within the internal combinatorial perturbation assessment, the MLTD-predicted perturbation space recovered structured pairwise response patterns among LAMB1 and its associated regulators (Fig. 4i,j). Removing field/flux information weakened and spatially dispersed the inferred knockout response, whereas expression correlation yielded substantially lower cell-level responses (Extended Data Fig. 8e). Together, these results show that MLTD progressively restores biologically expected but observationally obscured dependencies and reorganizes them into a selective, dataset-specific perturbation space.

### 3.5 MLTD resolves SOX2-centred perturbation programmes in head and neck cancer

SOX2 is a lineage-associated transcription factor implicated in squamous-cell identity, tumour-cell plasticity and stem-like programmes in head and neck squamous cell carcinoma^72–74^. We therefore applied MLTD to an HNSCC single-cell dataset spanning malignant, immune and stromal populations (Fig. 5a). SOX2 expression increased from normal and precancerous epithelial cells to primary tumour cells (Fig. 5b), but remained heterogeneous within the malignant compartment (Extended Data Fig. 9e). MLTD reconstructed a SOX2-centred dependency network linking SOX2 to transcriptional regulators and downstream response genes, including MYC-associated, epithelial-adhesion and stress-response components (Fig. 5c).

**Fig. 5.**
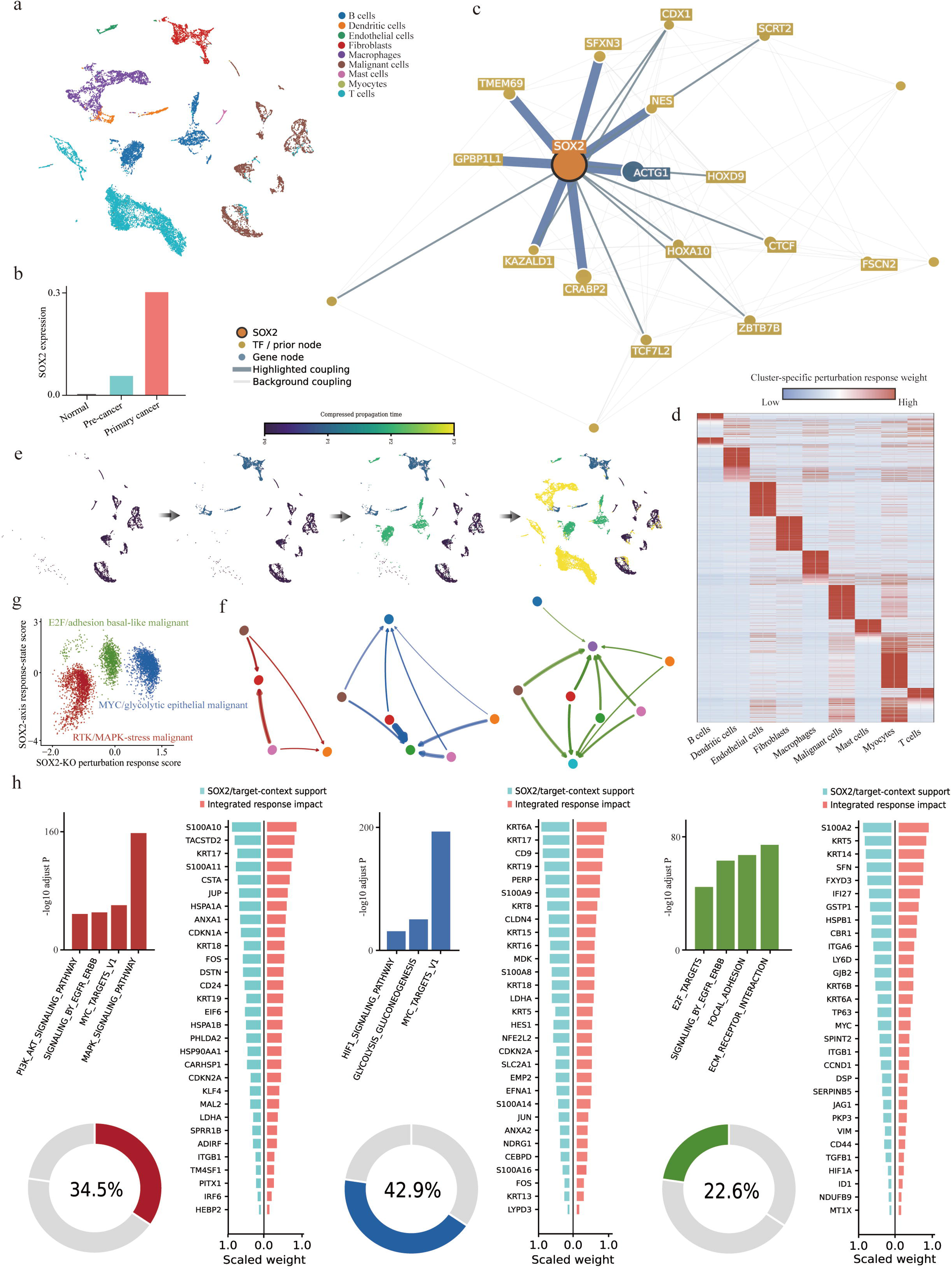
MLTD resolves SOX2-centred perturbation programmes across malignant cell states in head and neck cancer. **a,** UMAP representation of the HNSCC single-cell dataset, coloured by major cell type. **b,** Mean SOX2 expression in normal, precancerous and primary-cancer epithelial cells. **c,** SOX2-centred dependency network inferred by MLTD. The orange node denotes SOX2, gold nodes denote TF or prior-supported regulators and blue nodes denote response genes. Edge width indicates coupling strength; darker edges highlight the strongest inferred dependencies. **d,** Cluster-specific perturbation-response weights across the indicated cell populations. Rows represent response features and columns represent cells ordered by cell type; colour denotes the relative inferred response weight. **e,** Cell-level progression of the SOX2-associated response across the cellular manifold. Successive projections show the inferred propagation of response states; colour denotes compressed propagation time. **f,** State-specific communication networks associated with the three major SOX2-responsive malignant programmes. Nodes represent cell states and directed edges denote inferred communication strength. **g,** SOX2 target-response scores separating three malignant programmes: an E2F/adhesion-associated basal-like state, a MYC/glycolytic state and an RTK–MAPK stress-associated state. **h,** Pathway and gene-level decomposition of the three SOX2-responsive programmes. Left, pathway significance for each programme. Centre and right, relative SOX2/target-context support and integrated response impact for representative genes. Donut plots show the proportion of the integrated programme attributable to the highlighted response component.

The predicted SOX2 response was strongly cell-state dependent. Cluster-specific response weights formed discrete blocks across malignant and non-malignant populations (Fig. 5d), and projection along the inferred propagation trajectory revealed sequential engagement of distinct tumour-cell states (Fig. 5e). These states separated into E2F/adhesion-associated basal-like, MYC/glycolytic and RTK–MAPK stress-associated programmes (Fig. 5g). Such heterogeneity is consistent with the broader view that oncogenic transcription-factor activity is conditioned by lineage state and local regulatory context rather than expressed uniformly across tumour cells^75,76^. Their communication structures also differed, indicating that SOX2-associated responses propagated through state-specific intercellular routes (Fig. 5f).

Pathway-level decomposition showed that each SOX2-responsive state was supported by a distinct combination of target-context evidence and integrated perturbation impact (Fig. 5h). Basal-like cells were dominated by MAPK- and adhesion-related programmes, MYC-high malignant cells by glycolytic and proliferative programmes, and stress-associated cells by RTK, focal-adhesion and receptor-interaction pathways. SOX2 and MYC have been reported to cooperate in maintaining proliferative and undifferentiated tumour states^77,78^, and the recovery of a MYC-associated response axis therefore supports a context-specific regulatory connection rather than uniform SOX2–MYC co-expression across all cells.

MLTD also outperformed CPA^79^, scGPT^44^ and GEARS^19^ in recovering perturbation-associated expression trends (Extended Data Fig. 9a). Same-trend recovery remained high across multiple tumour datasets and increased with target-gene connectivity (Extended Data Fig. 9b,c). Five-stage ablation showed that the full response required prior, representation, communication, pseudotime and perturbation-flux components, with no single layer reproducing the full support score (Extended Data Fig. 9d). Independent expression maps and marker profiles further confirmed that the inferred programmes occupied distinct cellular regions rather than merely reflecting SOX2 abundance (Extended Data Fig. 9f,g). Together, these results show that MLTD resolves a heterogeneous, model-inferred SOX2-centred perturbation architecture in HNSCC, linking target-specific regulatory support to distinct malignant-state programmes and state-specific communication patterns.

### 3.6. MLTD reveals tumour–endothelial coupling associated with EndMT in renal cancer

Endothelial plasticity and endothelial-to-mesenchymal transition (EndMT) contribute to vascular remodelling, stromal activation and therapy resistance in renal tumours^80–82^. We therefore applied MLTD to renal cancer datasets containing multiple RCC subtypes and matched vascular and stromal populations. MLTD preserved the major malignant and non-malignant compartments while resolving an endothelial– pericyte axis that was only weakly represented in the original expression manifold (Fig. 6a,b). The reconstructed dependency space integrated expression, differential, network and communication support, with MLTD-specific evidence accounting for a substantial fraction of the final tumour-associated coupling (Fig. 6c).

**Fig. 6.**
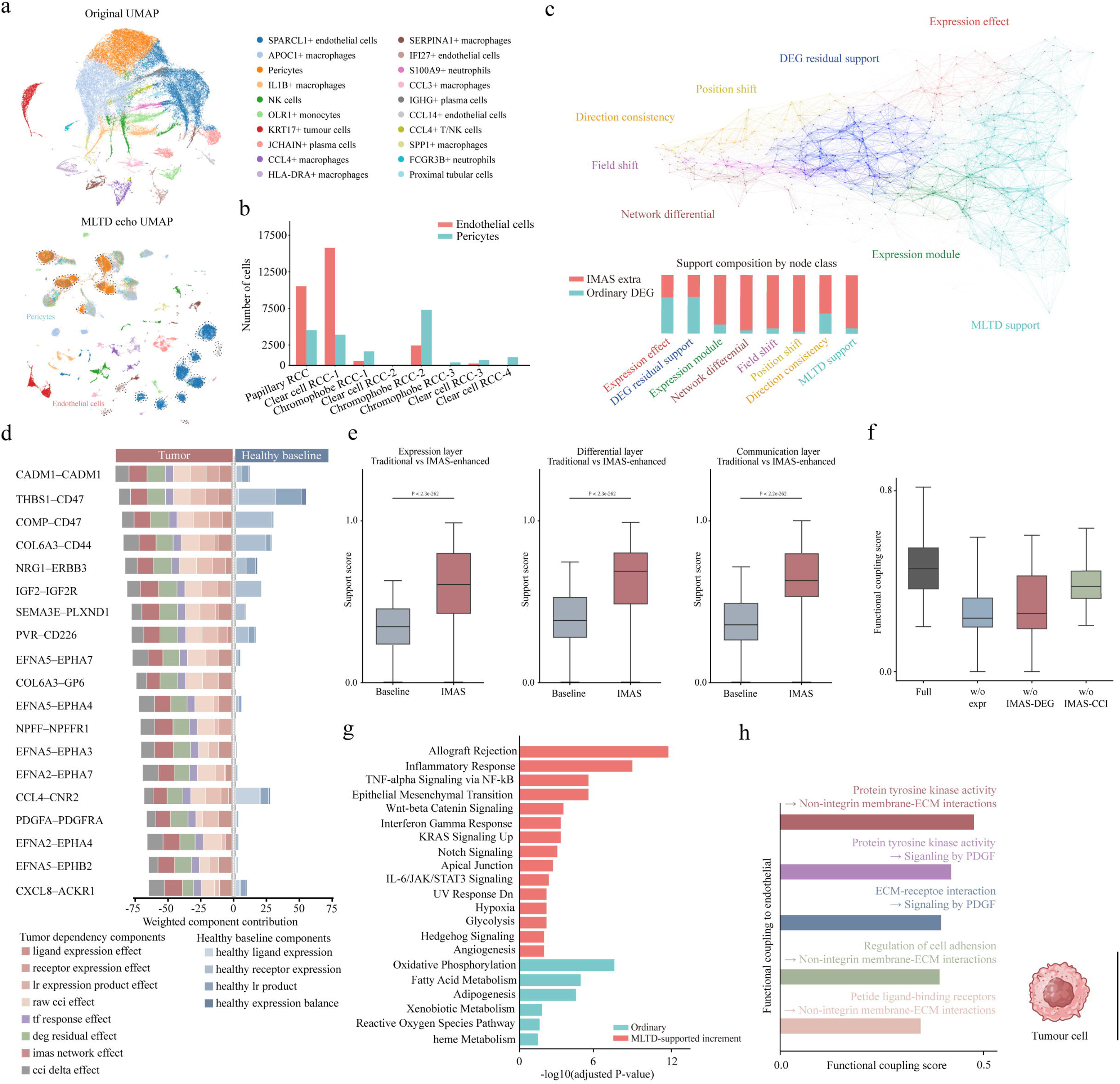
MLTD resolves tumour–vascular coupling associated with endothelial plasticity in renal cancer. **a,** Original and MLTD echo UMAP representations of the renal cancer single-cell atlas. Cells in the original UMAP are coloured by annotated cell type; the MLTD echo UMAP highlights target-relevant endothelial and pericyte compartments recovered in the dependency space. **b,** Numbers of endothelial and pericyte cells across the indicated renal cancer subtypes. Colours denote endothelial cells and pericytes. **c,** Integrated dependency network and support composition underlying the inferred tumour–vascular programme. Top, network representation of expression effect, DEG residual support, expression modules, network differential, positional and directional shifts, and MLTD support. Bottom, fraction of support contributed by ordinary differential-expression evidence and MLTD-specific increments for each node class. **d,** Weighted ligand– receptor contributions in tumour and healthy-baseline communication. Rows denote representative ligand–receptor pairs; stacked segments indicate the contributions of ligand expression, receptor expression, ligand–receptor product, raw CCI effect, TF-response effect, DEG residual effect, MLTD network effect and cell-state differential. **e,** Support scores for tumour–vascular coupling in the expression, differential and communication layers, comparing the conventional baseline with MLTD-enhanced inference. Boxes show the median and interquartile range; whiskers indicate the plotted range. P values are shown above each comparison. **f,** Functional coupling scores for the full model and models lacking expression-derived, MLTD-DEG or MLTD-CCI support. Boxes show the median and interquartile range. **g,** Enriched pathways associated with tumour–vascular coupling. Bars show adjusted significance for pathways detected by conventional analysis and the additional MLTD-supported increment. **h,** Representative functional couplings from tumour-cell programmes to endothelial responses. Bar length denotes coupling strength; labels indicate the inferred upstream activity and downstream endothelial pathway.

This coupling was concentrated in tumour–vascular interactions. Compared with the healthy baseline, tumour samples showed redistributed ligand–receptor support involving CADM1–CADM1, THBS1–CD47, collagen–CD44, NRG1–ERBB3, IGF2–IGF2R and multiple Ephrin–EPH interactions (Fig. 6d). Incorporating MLTD increased support across expression, differential and communication layers relative to conventional analysis, whereas removal of MLTD-derived expression or CCI evidence reduced the functional coupling score (Fig. 6e,f). These gains preferentially recovered inflammatory, TNF–NF-κB, epithelial–mesenchymal transition, WNT, JAK–STAT, hypoxia and angiogenic programmes (Fig. 6g), pathways known to coordinate tumour– vascular remodelling and mesenchymal activation^83–85^. The resulting model linked tumour-cell receptor-tyrosine-kinase and extracellular-matrix activity to endothelial adhesion and PDGF-associated signalling (Fig. 6h).

The inferred structure was not explained by generic imputation or differential expression alone. MLTD-supported genes occupied a restricted subset of the tumour-versus-normal differential landscape, and MLTD achieved higher overall mechanism scores than GRN-directed virtual knockout, GNN perturbation, latent-embedding perturbation or linear baselines (Extended Data Fig. 10a–c). Unlike MAGIC or ALRA, which largely reconstructed the existing differential-expression core, MLTD retained the raw core while adding context-supported genes that were not prominent in the original matrix (Extended Data Fig. 10d). Thus, the inferred tumour–endothelial programme reflected mechanistic augmentation rather than uniform smoothing of sparse expression.

We next evaluated this prediction in an independent spatial dataset. Xenium profiling identified activated or venous, blood and lymphatic endothelial populations together with pericytes, fibroblasts and tumour epithelial cells (Extended Data Fig. 11a,b). Within vascular regions, endothelial markers ERG and PECAM1 spatially overlapped with localized ACTA2-and VIM-high domains, accompanied by ANGPT2, ADGRL4 and VWF expression (Extended Data Fig. 11c). These cells showed subtype-specific mesenchymal, adhesion, cytoskeletal and vascular-abnormality programmes, with enrichment of focal adhesion, integrin, PI3K–Akt, smooth-muscle contraction and cell-adhesion pathways (Extended Data Fig. 11d,e). Together, these data support an MLTD-predicted tumour–endothelial coupling programme associated with spatially restricted EndMT-like states in renal cancer.

## Discussion

Tumour single-cell multiomic datasets provide direct access to transcriptional and chromatin-regulatory states, but their sparsity, heterogeneity and limited sample size make exhaustive reconstruction of tumour regulatory systems unrealistic from sequencing data alone^11,86,87^. IMAS was therefore designed around a narrower objective: to recover compact, target-specific regulatory architectures that remain traceable across molecular layers and can be prioritized for experimental testing. In this framework, IMAS denotes the pan-cancer foundation and target-adaptation system, whereas multi-layer target dependencies (MLTDs) are the target-aligned architectures used for downstream perturbation and mechanism discovery.

The central contribution of IMAS is the use of MLTDs as an intermediate representation between raw molecular measurements and mechanistic hypotheses. Existing methods typically infer transcriptional states, regulatory networks, cell–cell communication and perturbation responses as separate outputs^88–91^. IMAS instead asks whether a candidate signal remains supported across matched multiomic priors, target-adapted RNA–TF–regulatory-element relationships, communication context and perturbation-sensitive local structure. A signal is therefore not prioritized merely because it is differentially expressed, assigned a high model score or represented in a ligand–receptor catalogue. This does not establish a complete causal map, but narrows the hypothesis space to internally consistent and target-supported regulatory architectures.

Matched multiomic-only foundation learning was important for preserving cross-layer traceability. Large scRNA-seq collections provide broader cellular coverage, but do not directly observe regulatory-element usage in the same cells^92,93^. In our analyses, increasing RNA-only representation did not compensate for dilution of matched RNA– chromatin supervision: dependency support, regulatory-element traceability, dependency-core preservation and response anchoring all declined as the matched multiomic contribution was reduced. The value of the foundation corpus therefore lies not only in predictive scale, but in preserving transferable RNA–TF–regulatory-element structure before adaptation to a target dataset.

Target-aware adaptation then converted these transferable relations into dataset-specific MLTDs. In colorectal cancer, adapted dependencies differed substantially from native expression, conventional co-expression and the inherited pan-cancer hierarchy. Counterfactual deletion, rationale retention and layer-specific ablation further showed that the selected structures were compact and model-intervention sensitive. These results suggest that adaptation in IMAS acts less as generic batch correction and more as a mechanism-selection process that re-ranks inherited support according to the target cellular context.

MLTDs also provided a common structure through which perturbation-associated information could be propagated. Quantitative RNA and TF recovery was connected to RNA–TF–RE bridges, which in turn informed ligand–receptor communication and receiver-side TF responses. The resulting GNN–LTC ordering should not be interpreted as developmental or transcriptomic pseudotime. Instead, it represents a model-derived ordering of sender activity, receiver response and downstream regulatory context. This distinction is important because transcriptomic similarity alone does not necessarily encode how signalling is transmitted between cellular states^94,95^. The selective increase in TF-aware temporal alignment, despite limited changes in global forward-direction metrics, further indicates that MLTD primarily strengthened the coupling between communication and receiver regulation rather than uniformly increasing network directionality.

The LAMB1 analysis illustrates a second role of MLTD recovery: restoring biologically expected dependencies that are weakly represented in the observed matrix. Although LAMB1 is intrinsically linked to basement-membrane and extracellular-matrix biology^96,97^, ECM–receptor interaction was not prominent in the initial dependency field. Its support emerged progressively as representation, communication, temporal and perturbation-flux constraints were introduced. At the same time, the top-response hierarchy was repeatedly reorganized. This pattern argues against indiscriminate amplification of prior knowledge and instead supports selective recovery of target-supported relationships obscured by sparsity, low expression or indirect regulation.

The HNSCC and renal cancer analyses extended these observations to independent biological settings. In HNSCC, SOX2-centred MLTDs separated malignant cells into distinct basal-like, MYC-associated and RTK–MAPK stress-response programmes, indicating that perturbation response was conditioned by cell state rather than uniformly determined by SOX2 abundance. In renal cancer, MLTD-guided analysis preserved the ordinary differential-expression core while adding network-supported tumour–vascular dependencies. Spatial Xenium data subsequently identified endothelial regions with mesenchymal, adhesion and vascular-abnormality features, supporting the plausibility of the predicted tumour–vascular remodelling programme. These findings do not establish a direct tumour-RTK-to-EndMT mechanism, but they define a spatially supported hypothesis for focused experimental testing.

IMAS differs in this respect from imputation or complete matrix-reconstruction workflows. It does not replace the measured expression matrix or assume that unobserved regulatory states can be exhaustively reconstructed. Instead, it adds an MLTD-supported layer that can be projected back onto the native data while retaining the original cellular measurements. Augmentation therefore refers to the addition of traceable dependency support rather than substitution of the observed molecular state. Several limitations remain. IMAS infers regulatory and communication constraints from matched multiomic priors, transcriptomic context and ligand–receptor evidence rather than directly measured signalling events. Virtual perturbation remains a model-based prioritization strategy and cannot substitute for genetic or pharmacological intervention. The current analyses span several tumour types, but broader testing across additional diseases, non-neoplastic systems and prospective cohorts will be required to establish generality. Finally, spatial correspondence supports tissue-level plausibility but does not prove individual ligand–receptor, TF or regulatory-element mechanisms.

In summary, IMAS provides a pan-cancer multiomic foundation and adaptation framework, whereas MLTDs provide the target-specific representation through which perturbation, cross-layer regulation and intercellular communication are interpreted. By preserving native tumour states while adding compact and traceable dependency support, IMAS shifts mechanism discovery away from exhaustive network reconstruction and towards structured, experimentally actionable hypothesis prioritization.

## Data availability

The raw and processed single-cell RNA-sequencing data generated from the eight patient-derived renal tumours will be deposited in GEO under accession. Publicly available datasets analysed in this study were obtained from the Gene Expression Omnibus under accession codes GSE294559, GSE210566, GSE307240 and GSE181919. The ccRCC Xenium dataset was obtained from the 10x Genomics datasets portal under the title “Xenium In Situ Gene and Protein Expression data for FFPE Human Renal Cell Carcinoma”. Source data underlying the main and supplementary figures are provided with the paper. Processed matrices, cell annotations and model-derived analysis outputs will be made available through the associated data repository. Code and analysis scripts will be publicly available on GitHub at upon publication.

## Acknowledgements

We thank Prof. Yasusei Kudo (Tokushima University) and Dr Yuki Tobisawa (Gifu University) for insightful discussions and valuable advice.

## Author Contributions

D. Wu: Conceptualization; Methodology; Investigation; Data curation; Visualization; Writing – original draft. T. Inubushi: Conceptualization; Data curation; Writing – review & editing; Supervision; Funding acquisition. Both authors contributed to interpretation of the results and approved the final manuscript. T. Yamashiro: Supervision. Writing – review & editing.

## Appendix

**Table S1.**
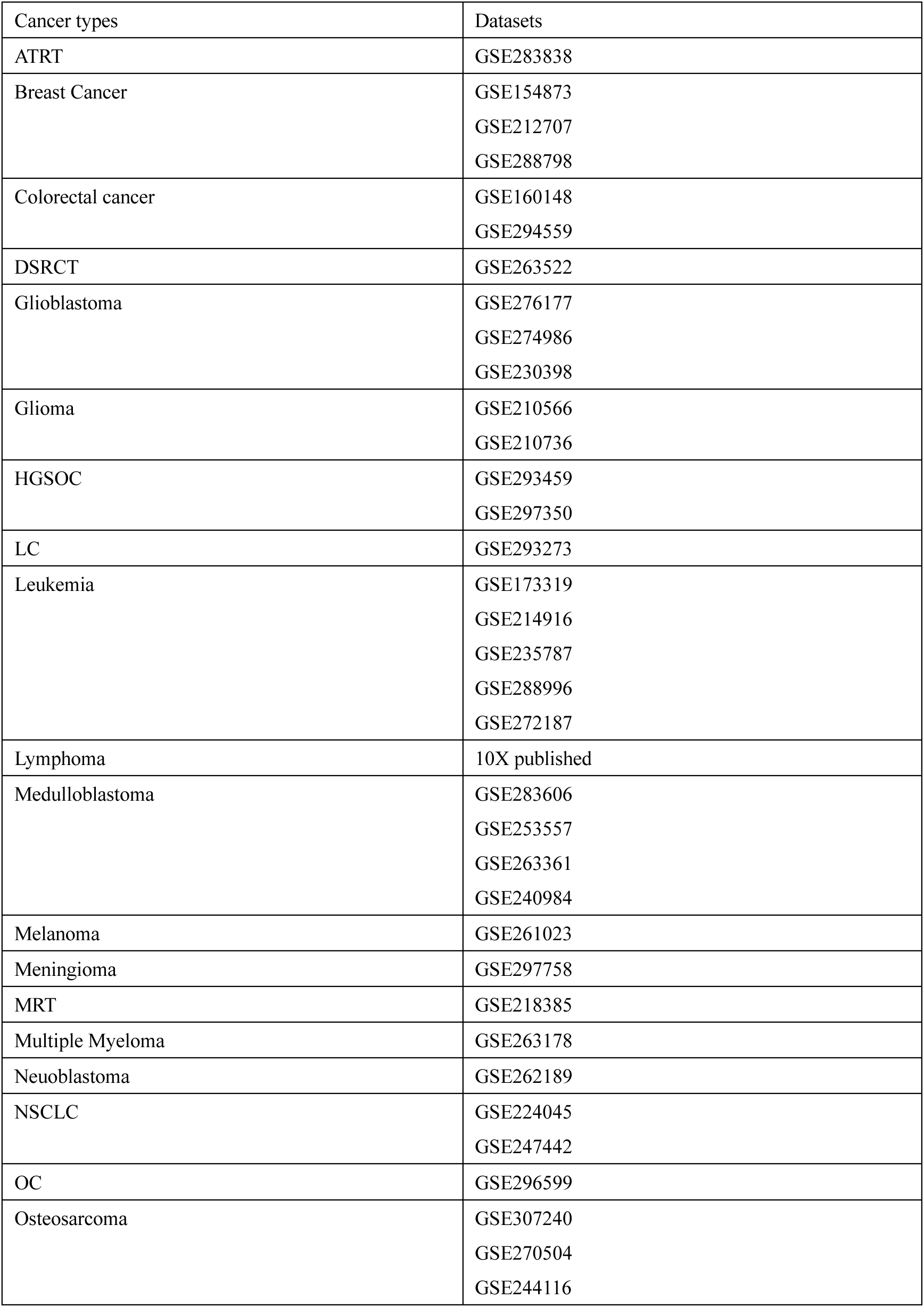
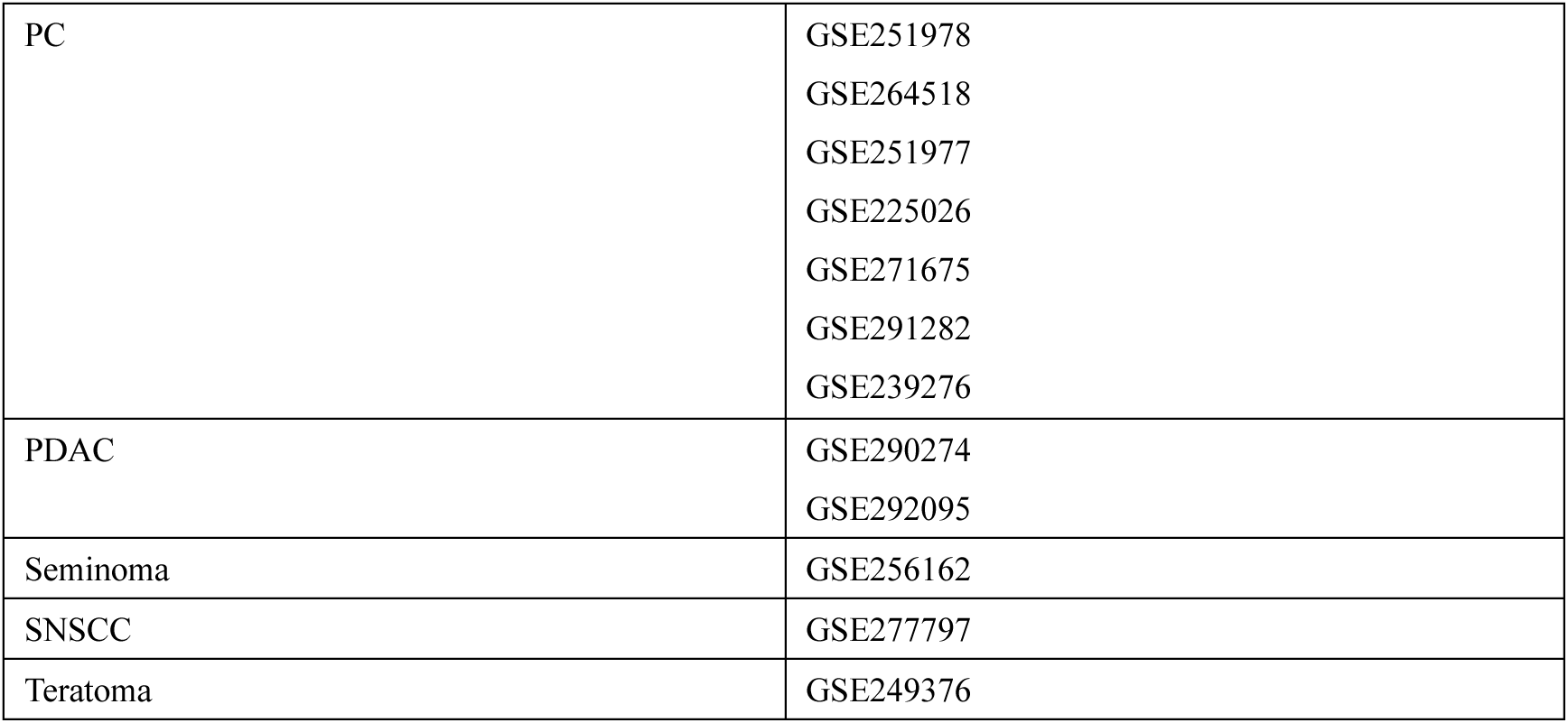
datasets for pan-cancer model.

**Table S2.** datasets for independent validation.

| Cancer types | Datasets | Knockout gene |
| --- | --- | --- |
| Breast Cancer | GSE245601 | ESR1 |
|  | GSE145326 | FGFR |
| CRC | GSE232525 | VEGFB |
| Endometrial carcinoma | GSE225689 | NTN1 |
| HNSCC | GSE173647 | EGFR |
|  | GSE137524 | EGFR |
| Melanoma | GSE229908 | BRAF |
|  | GSE164897 | BRAF |
| PC | GSE150949 | EGFR |

**Extend Data Fig. 1.**
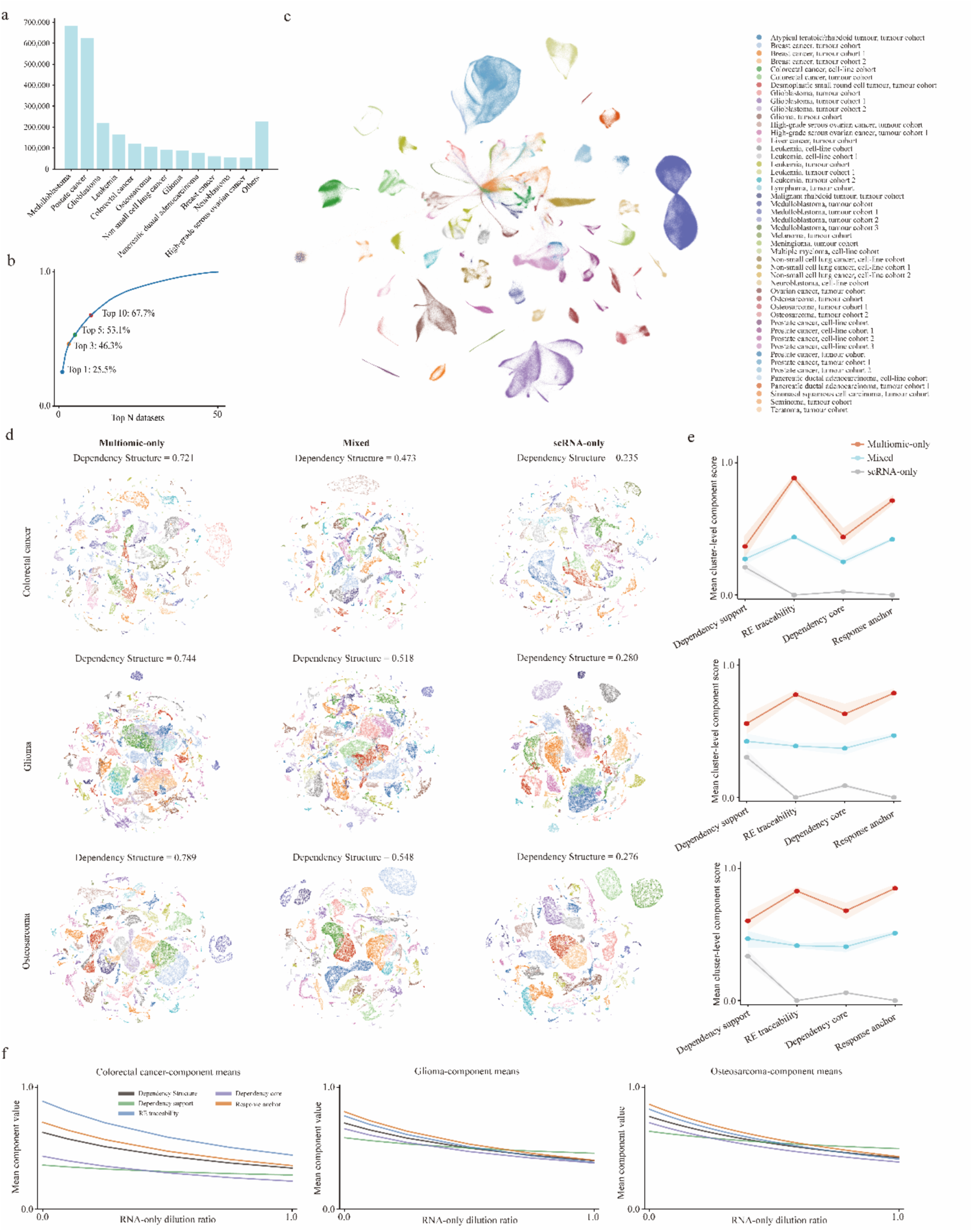
Matched multiomic supervision preserves cross-layer information required for MLTD recovery. **a,** Numbers of matched single cells contributed by the indicated tumour types in the pan-cancer foundation corpus. **b,** Cumulative fraction of cells contributed by the top-ranked datasets. Labels indicate the fractions contributed by the top 1, 3, 5 and 10 datasets. **c,** Joint embedding of the matched pan-cancer multiomic corpus. Cells are coloured by tumour type, cohort and experimental source as indicated. **d,** Target-cell dependency structures inferred in colorectal cancer, glioma and osteosarcoma using multiomic-only, mixed multiomic–scRNA and scRNA-only foundation settings. Cells are coloured by target cell state. The dependency-structure score is shown above each embedding. **e,** Mean cluster-level component scores for dependency support, regulatory-element traceability, dependency-core preservation and response anchoring under multiomic-only, mixed and scRNA-only training. Panels correspond to colorectal cancer, glioma and osteosarcoma. Lines show mean values and shaded regions indicate variability across clusters. **f,** Mean component values as matched multiomic information was progressively diluted with RNA-only data. Curves show dependency structure, dependency support, regulatory-element traceability, dependency-core preservation and response anchoring in colorectal cancer, glioma and osteosarcoma. RNA-only dilution progressively weakens cross-layer support

**Extend Data Fig. 2.**
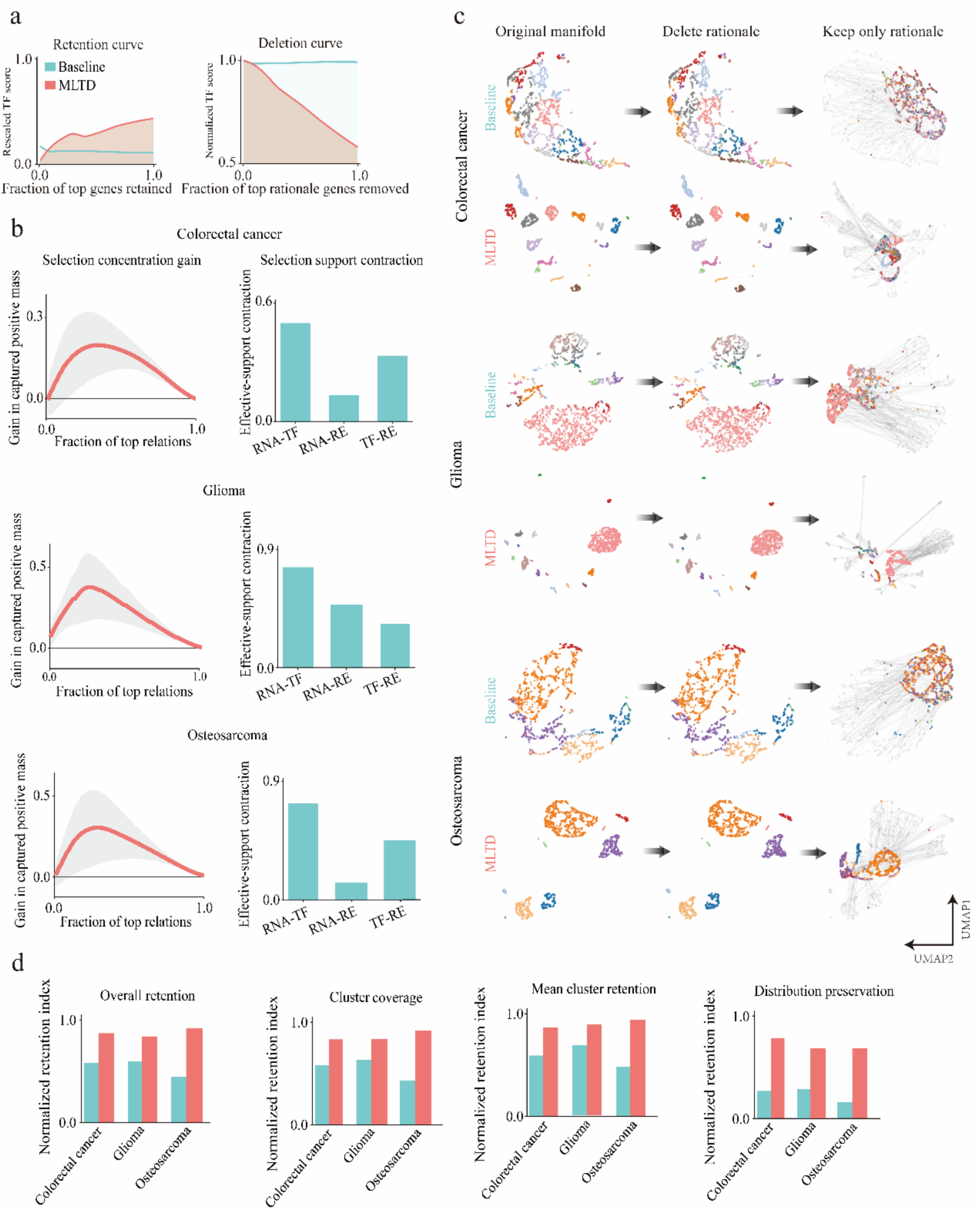
MLTD-prioritized relations preserve molecular support and cellular-state structure across tumour types. **a,** Retention and deletion curves for MLTD-prioritized relations. Left, rescaled TF support as an increasing fraction of top-ranked rational features was retained. Right, normalized TF score as an increasing fraction of rationale genes was removed. MLTD is compared with the corresponding baseline selection. **b,** Selection concentration and effective-support contraction in colorectal cancer, glioma and osteosarcoma. Left, gain in captured positive mass as a function of the fraction of top-ranked relations. Lines show the mean and shaded regions indicate variability across repetitions. Right, effective-support contraction across RNA–TF, RNA–RE and TF–RE relation classes. **c,** Manifold-level rationale-retention analysis in colorectal cancer, glioma and osteosarcoma. For each tumour type, the original manifold, the manifold after deletion of the selected rationale and the structure retained when only the rationale was preserved are shown for the baseline and MLTD selections. Grey lines indicate retained neighbourhood relations. **d,** Quantitative comparison of overall retention, cluster coverage, mean cluster retention and distribution preservation between baseline and MLTD across the three tumour types. Values were normalized within each metric.

**Extend Data Fig. 3.**
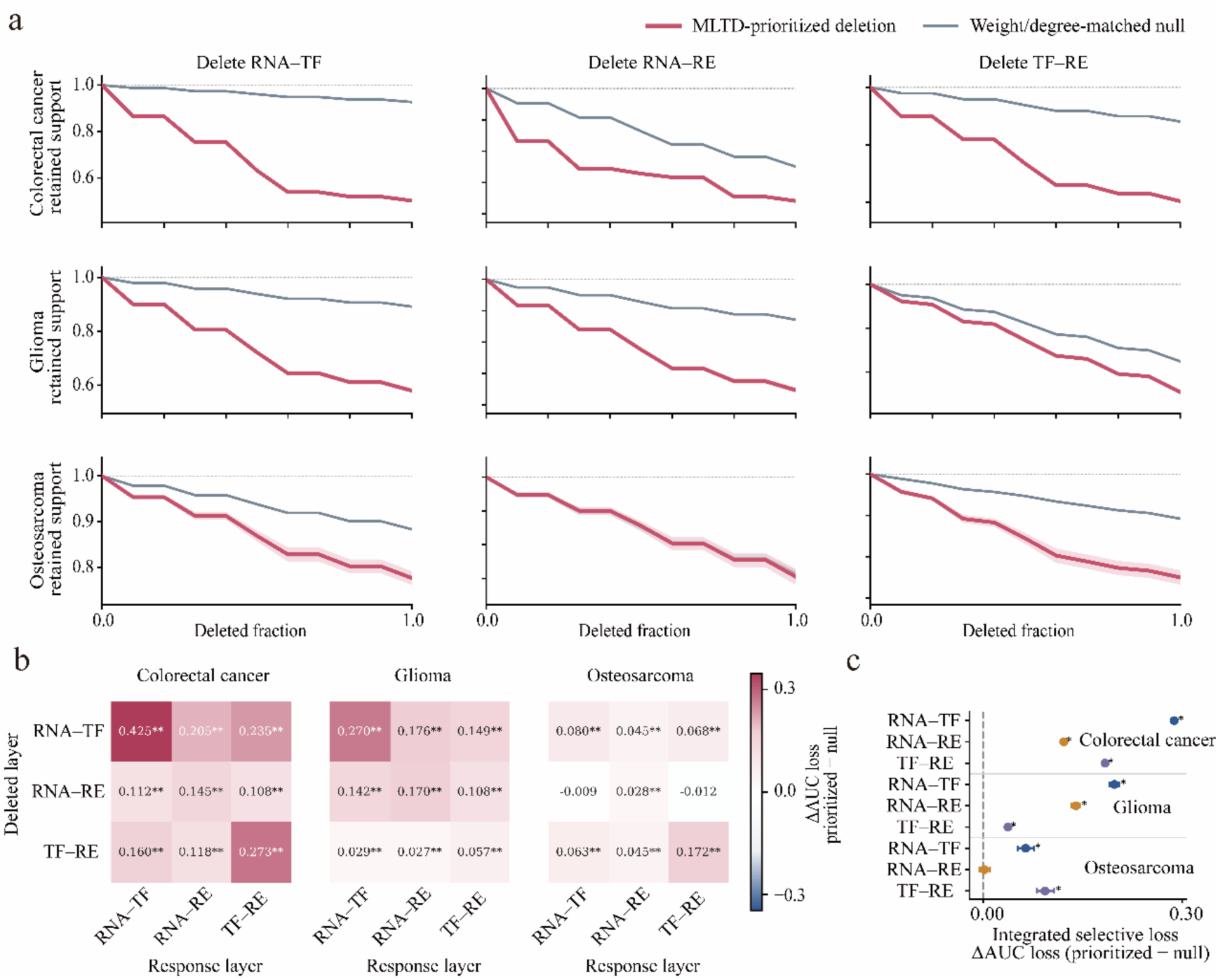
Layer-specific deletion reveals coordinated RNA–TF, RNA– RE and TF–RE dependency. **a,** Retained support after progressive deletion of RNA–TF, RNA–RE or TF–RE relations in colorectal cancer, glioma and osteosarcoma. Red lines indicate MLTD-prioritized deletion and grey lines indicate weight-and degree-matched null deletion. Shaded regions show variability across repeated deletions where available. **b,** Changes in response-layer AUC after deleting each relation class. Rows indicate the deleted layer and columns indicate the response layer. Values show the AUC loss for MLTD-prioritized deletion relative to the matched null; asterisks indicate statistically significant differences. Positive values indicate greater loss after MLTD-prioritized deletion. **c,** Integrated selective loss across deleted relation classes and tumour types. Points show the difference in integrated AUC loss between prioritized and matched-null deletion; horizontal lines indicate uncertainty intervals. The dashed vertical line denotes no excess loss, and asterisks indicate significant effects.

**Extend Data Fig. 4.**
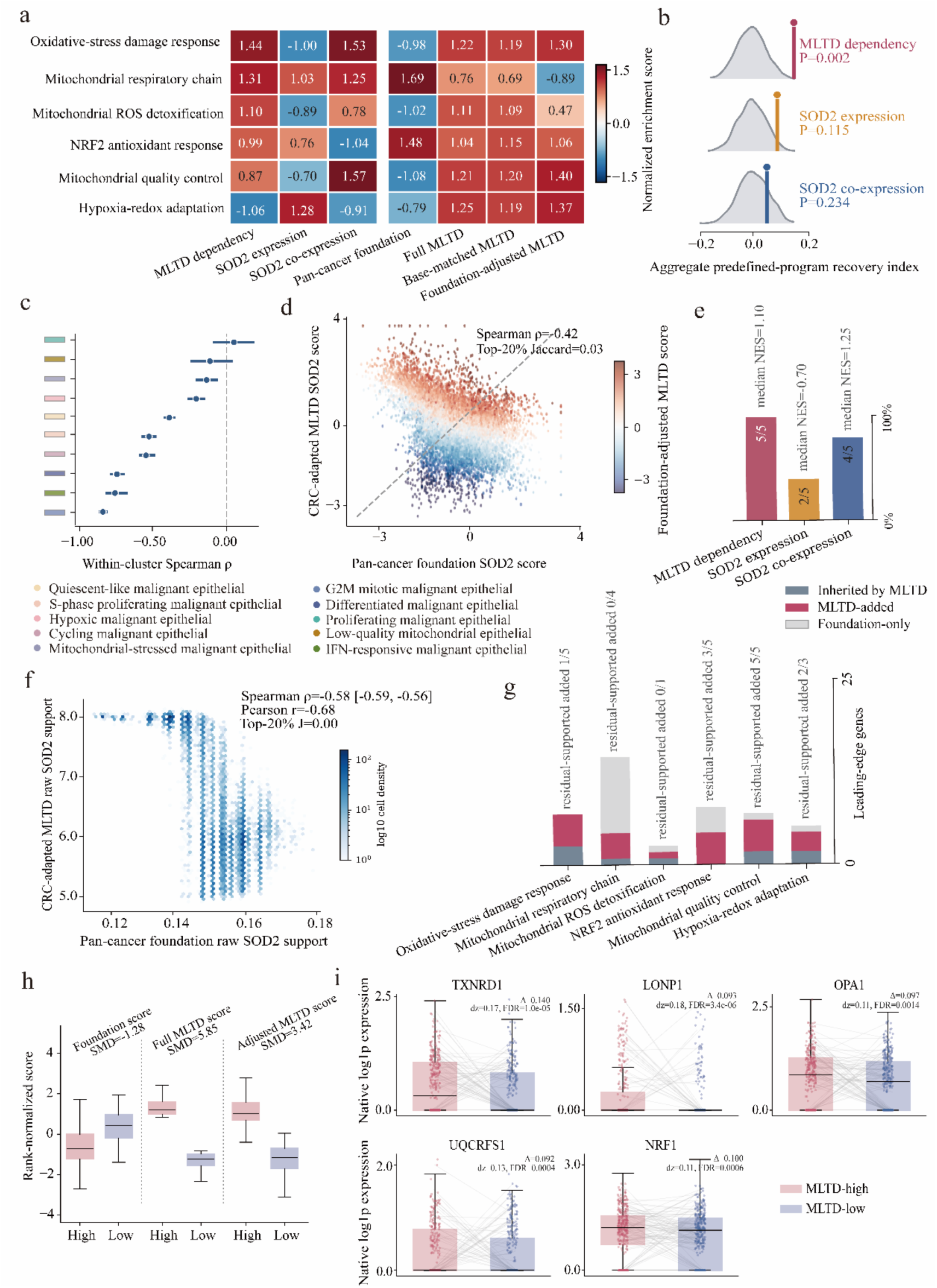
Target adaptation reorganizes the SOD2-associated dependency programme. **a,** Recovery scores of predefined mitochondrial and oxidative-stress programmes using MLTD dependency, native SOD2 expression, SOD2 co-expression, the pan-cancer foundation score, full MLTD, base-matched MLTD and foundation-adjusted MLTD. Values are normalized within programme. **b,** Null distributions of the aggregate predefined-program recovery index for MLTD dependency, native SOD2 expression and SOD2 co-expression. Vertical lines indicate observed values; empirical P values are shown. **c,** Within-cluster Spearman correlations between pan-cancer foundation and CRC-adapted MLTD SOD2 scores. Points show cluster estimates and horizontal lines indicate confidence intervals. Colours denote malignant epithelial cell states; the dashed line marks zero correlation. **d,** Cell-level comparison of pan-cancer foundation and CRC-adapted MLTD SOD2 scores. Points are coloured by foundation-adjusted MLTD score. The global Spearman correlation and top-20% Jaccard overlap are shown. **e,** Number of predefined programmes supported by MLTD dependency, native SOD2 expression and SOD2 co-expression. Bars show the number of programmes exceeding the normalized enrichment-score threshold. **f,** Relationship between pan-cancer foundation support and CRC-adapted MLTD raw SOD2 support. Point density is shown on a logarithmic scale; correlation statistics are indicated. **g,** Leading-edge genes contributed by inherited MLTD support, MLTD-added support and foundation-only support across predefined programmes. Numbers above bars indicate the corresponding gene counts. **h,** Rank-normalized foundation, full-MLTD and foundation-adjusted MLTD scores in MLTD-high and MLTD-low cells. SMD values are shown above each comparison. **i,** Native log-transformed expression of representative leading-edge genes in MLTD-high and MLTD-low cells. Points represent cells, boxes show the median and interquartile range, whiskers extend to 1.5 times the interquartile range and grey lines connect matched observations. Effect sizes and false-discovery rates are shown above each panel.

**Extend Data Fig. 5.**
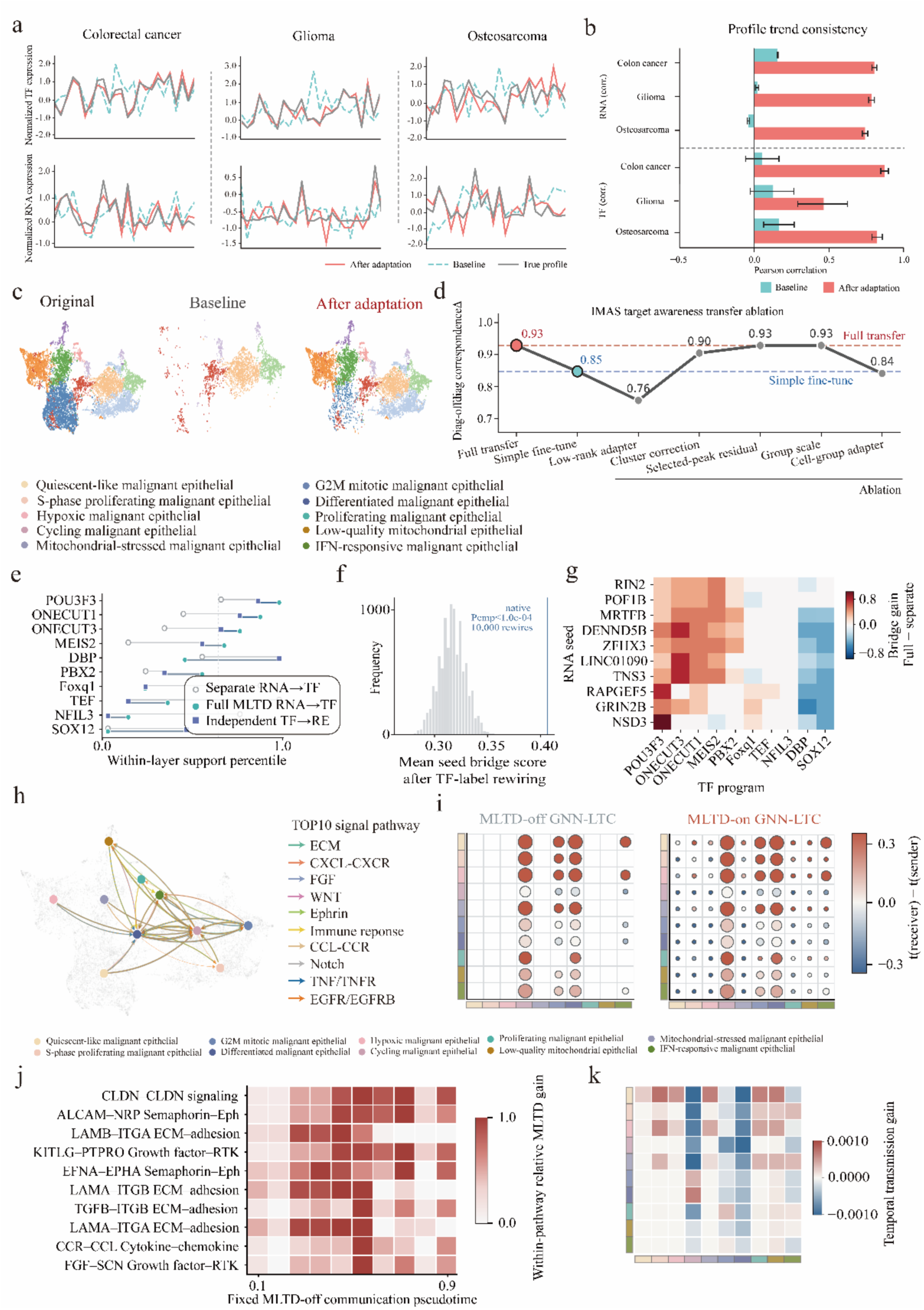
Target adaptation supports quantitative state recovery and cross-layer communication propagation. **a,** Representative normalized TF-expression and RNA-expression profiles in colorectal cancer, glioma and osteosarcoma. Red lines show profiles after adaptation, dashed turquoise lines show the baseline and grey lines show the observed profiles. **b,** Pearson correlations between inferred and observed RNA or TF profile trends before and after adaptation across the three tumour types. Bars show summary estimates and error bars indicate uncertainty. **c,** Cellular manifolds for the original data, baseline reconstruction and reconstruction after adaptation. Cells are coloured by malignant epithelial state. **d,** Target-awareness transfer ablation. Diagonal-to-off-diagonal correspondence is shown for full transfer, simple fine-tuning and the indicated component deletions. Dashed horizontal lines mark the full-transfer and simple-fine-tuning references. **e,** Within-layer support percentiles for representative TF programmes using separately inferred RNA–TF relations, full MLTD RNA–TF integration and independent TF–RE support. Higher values indicate stronger cluster-resolved matching between inferred and observed states. **f,** Null distribution of the mean RNA-seed bridge score after TF-label rewiring. The vertical line indicates the observed score and the empirical P value is shown. **g,** Gain in RNA-seed–TF bridge score produced by full MLTD relative to separate-layer inference. Rows denote RNA seeds and columns denote TF programmes. **h,** Cluster-level communication network formed by the ten principal signalling modules. Nodes denote malignant epithelial states, edge width denotes communication strength and edge colour indicates pathway class. **i,** MLTD-off and MLTD-on GNN–LTC communication matrices. Bubble size denotes temporally weighted communication strength and colour denotes the receiver–sender pseudotime difference. Marginal colour strips indicate cluster identity. **j,** Relative MLTD gain for representative ligand– receptor pathways across fixed MLTD-off communication pseudotime. **k,** Sender–receiver matrix of temporal-transmission gain. Colour denotes the difference in temporally weighted communication between MLTD-on and MLTD-off.

**Extend Data Fig. 6.**
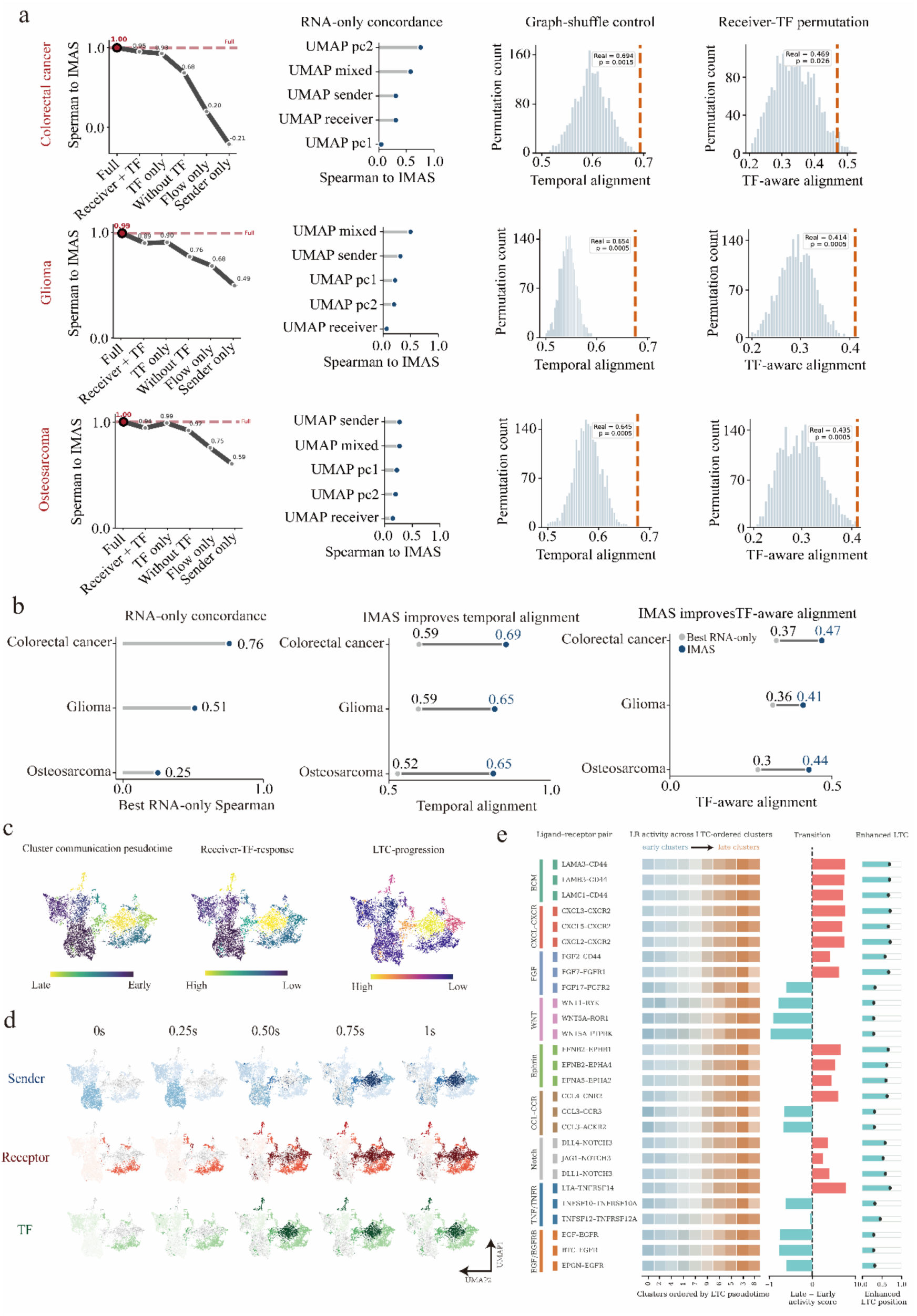
Controls and dynamic characterization of MLTD-informed communication pseudotime. **a,** Validation of communication pseudotime in colorectal cancer, glioma and osteosarcoma. For each tumour type, the panels show concordance of feature-ablation models with the full model, concordance with RNA-only manifold orderings, graph-shuffle null distributions for temporal alignment and receiver-TF permutation null distributions for TF-aware alignment. Dashed orange lines indicate observed values and empirical P values are shown. **b,** Cross-tumour summary of the best RNA-only concordance, temporal alignment and TF-aware temporal alignment. Grey points indicate the best RNA-only control and blue points indicate the full MLTD-informed model. **c,** Cell-level projections of cluster communication pseudotime, receiver-TF response and LTC progression on the cellular manifold. **d,** Dynamic sender, receptor and TF signals projected onto the manifold at the indicated LTC rollout times. Colour intensity denotes the corresponding signal magnitude. **e,** Activity of representative ligand–receptor pairs across clusters ordered by LTC pseudotime. Rows are grouped by signalling module. The adjacent columns show late-to-early transition scores and enhanced LTC positions.

**Extend Data Fig. 7.**
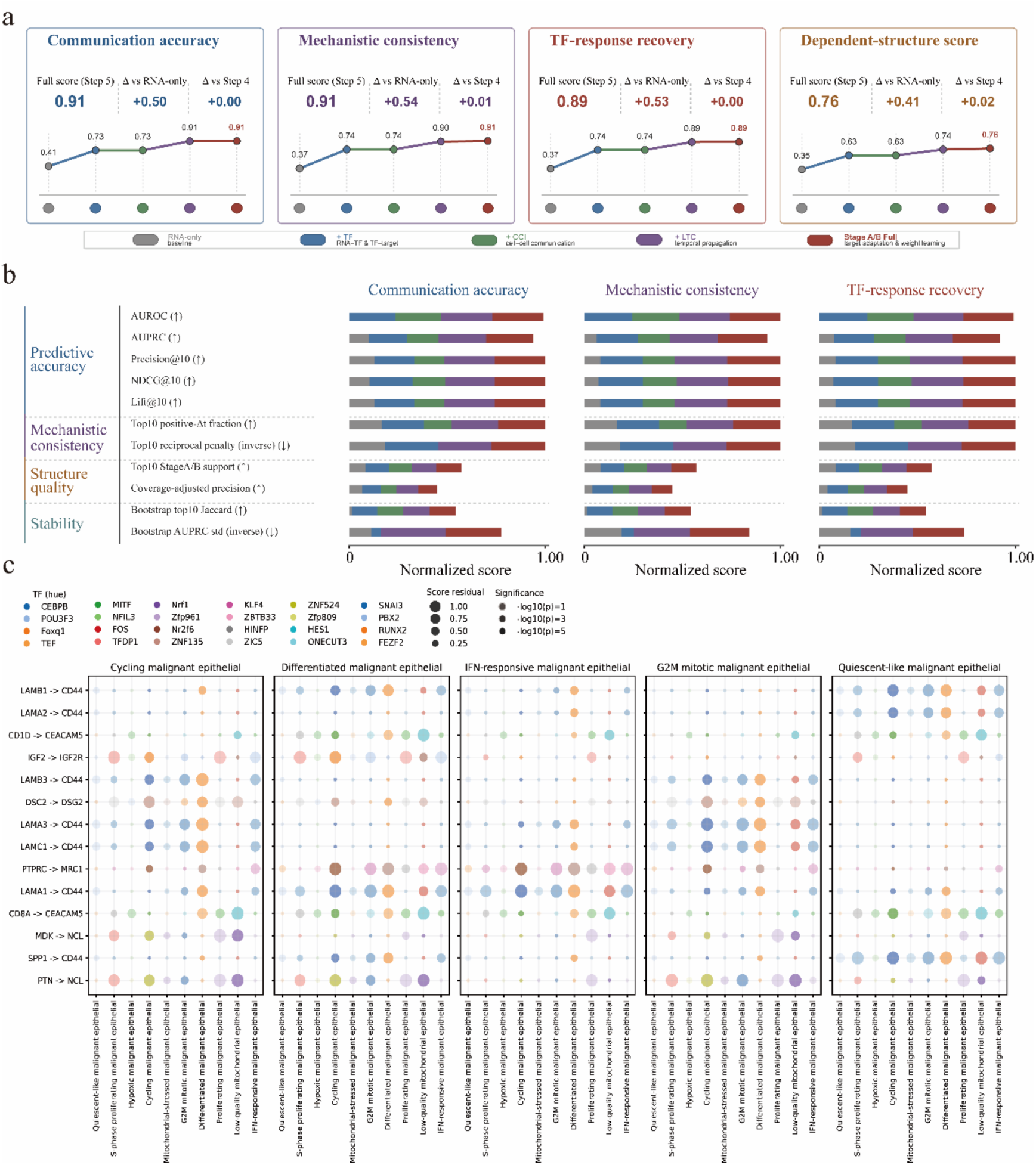
Stage-wise contributions to communication prediction and TF-aware propagation **a,** Stage-wise performance for communication accuracy, mechanistic consistency, TF-response recovery and dependency-structure score. Values are shown for the RNA-only baseline, addition of RNA–TF and TF-target information, addition of cell–cell communication, addition of LTC temporal propagation and the full MLTD model. Summary values report the full-model score and gains relative to the RNA-only baseline and the preceding stage. **b,** Normalized stage-wise contributions to individual metrics underlying communication accuracy, mechanistic consistency and TF-response recovery. Metrics are grouped into predictive accuracy, mechanistic consistency, structure quality and stability. Colours denote the model component added at each stage. **c,** TF-aware ligand–receptor structure across five malignant epithelial states. Rows denote ligand–receptor pairs and columns denote receiver-cell states within each sender-state panel. Bubble size represents the residual interaction score, colour denotes the associated TF programme and opacity or outline intensity reflects statistical significance.

**Extend Data Fig. 8.**
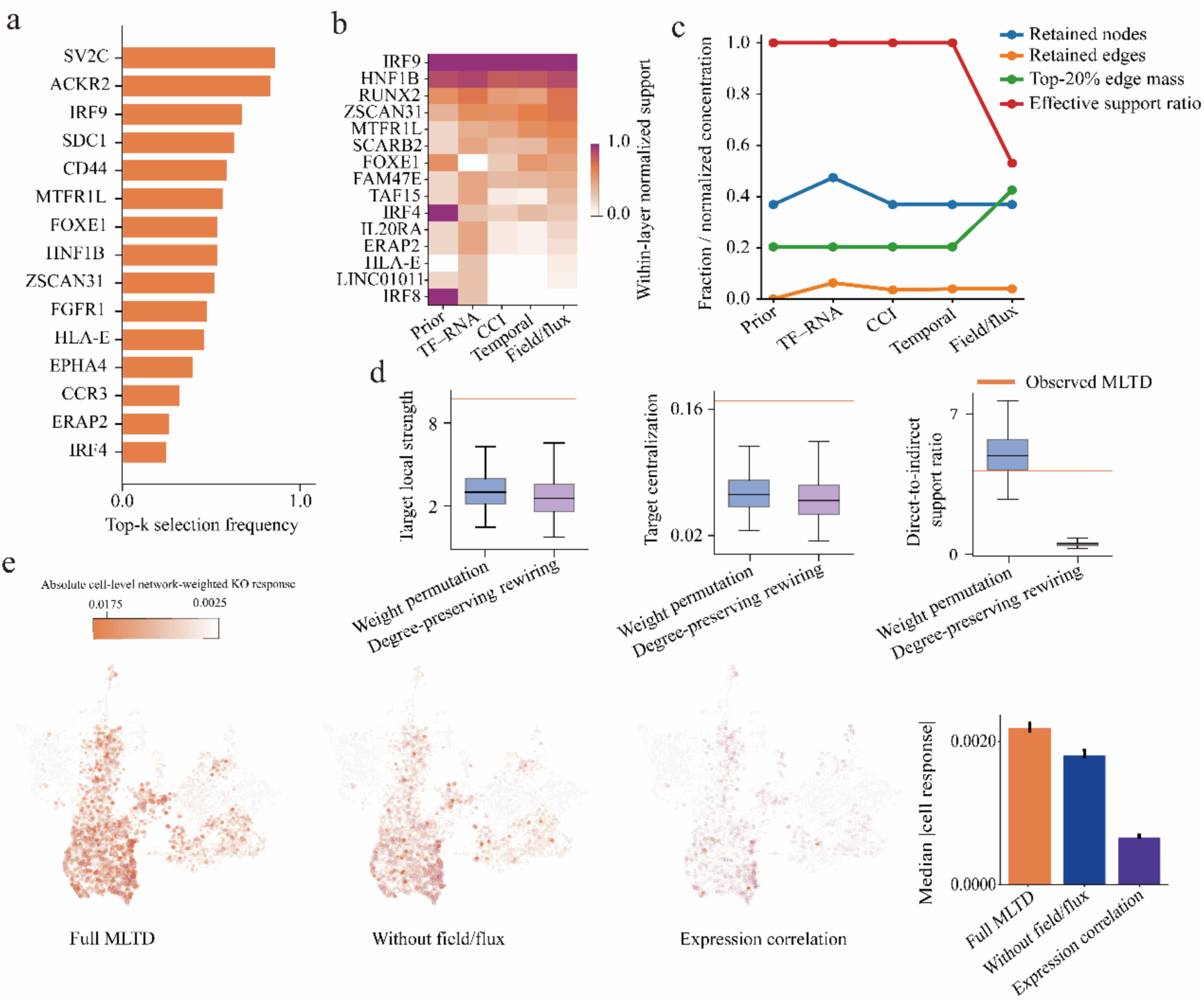
Structural concentration and perturbation-support analysis of the LAMB1 dependency field **a,** Frequency with which the indicated genes were selected among the top-ranked LAMB1 responders across model stages or repeated analyses. **b,** Within-layer normalized support for recurrent responders across the prior, TF–RNA, CCI, temporal and field/flux dependency layers. Rows denote candidate responders and columns denote successive sources of constraint. **c,** Structural concentration of the dependency field across successive stages. Lines show the fractions of retained nodes and retained edges, the mass contained in the top 20% of edges and the effective-support ratio, each normalized to the corresponding reference value. **d,** Network-control analysis comparing weight permutation and degree-preserving rewiring. Box plots show target-local strength, target centralization and the direct-to-indirect support ratio. Orange horizontal lines indicate the observed MLTD values. Boxes show the median and interquartile range; whiskers indicate the plotted range. **e,** Cell-level absolute network-weighted virtual knockout response under full MLTD, MLTD without field/flux information and an expression-correlation baseline. Manifold projections show the spatial distribution of predicted responses; the bar plot summarizes the median cell-level response, with error bars indicating uncertainty.

**Extend Data Fig. 9.**
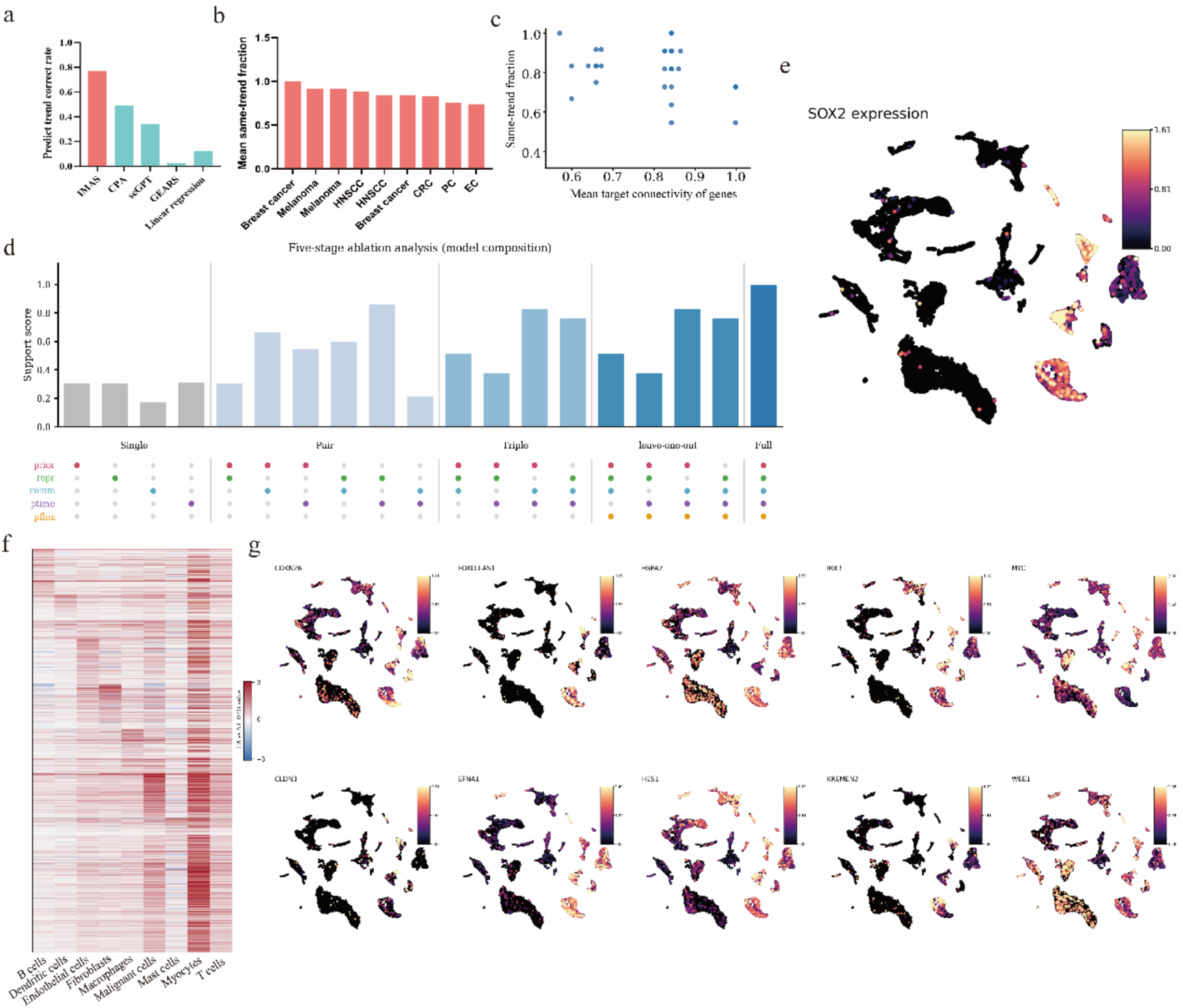
Benchmarking, ablation and expression context of SOX2 perturbation recovery **a,** Perturbation-trend recovery accuracy for MLTD, CPA, scGPT and GEARS. Bars show the fraction of genes with correctly predicted response direction. **b,** Mean same-trend fraction across the indicated cancer datasets. **c,** Relationship between mean target-gene connectivity and same-trend recovery across datasets. Each point represents one target–dataset evaluation. **d,** Five-stage ablation analysis of SOX2-response support. Bars show the support score obtained using single components, component pairs, component triples, leave-one-out models and the full model. The dot matrix indicates inclusion of prior, representation, communication, pseudotime and field/flux components. **e,** Cell-level SOX2 expression projected onto the HNSCC UMAP. **f,** Cell type-resolved expression heat map for SOX2-associated response genes. Values are scaled within gene. **g,** UMAP projections of representative genes associated with the inferred SOX2-responsive programmes, including CDKN2B, FOXD3-AS1, HSPA2, IRX3, MYC, CLDN3, EFNA1, HES1, KREMEN2 and WEE1. Colour denotes normalized expression.

**Extend Data Fig. 10.**
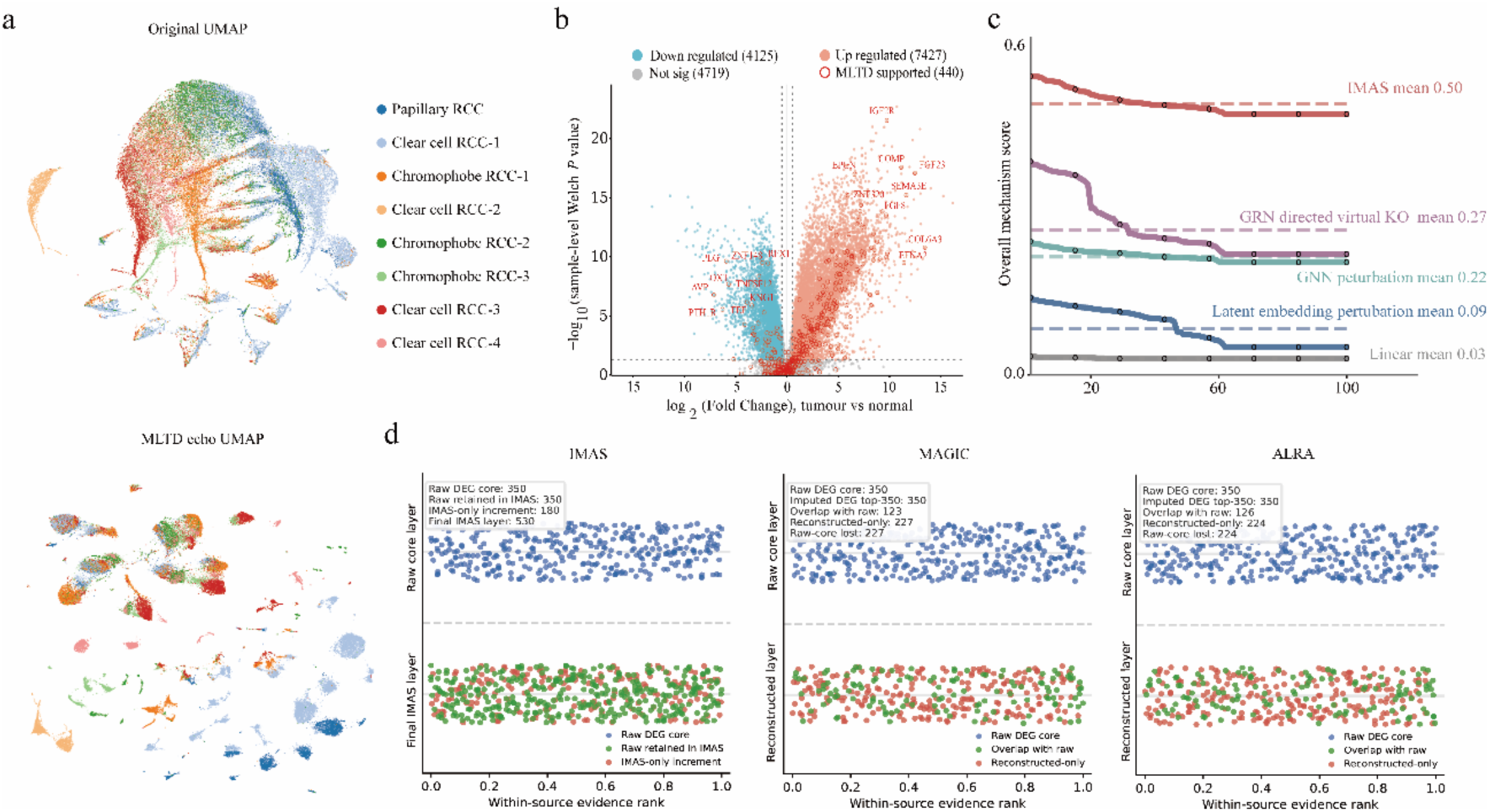
MLTD augments tumour-associated differential signals beyond imputation and generic perturbation baselines **a,** Original and MLTD echo UMAP representations of renal cancer cells, coloured by tumour subtype. **b,** Tumour-versus-normal differential-expression landscape. Points indicate downregulated, upregulated or non-significant genes; outlined points denote genes supported by MLTD. Selected genes are labelled. **c,** Overall mechanism scores across increasing evidence-rank thresholds for MLTD, GRN-directed virtual knockout, GNN perturbation, latent-embedding perturbation and a linear baseline. Dashed horizontal lines indicate mean scores for each method. **d,** Comparison of raw differential-expression cores and reconstructed layers generated by MLTD, MAGIC and ALRA. The upper panels show genes retained from the raw DEG core; the lower panels show genes retained or added in the final reconstructed layer. Colours distinguish raw-core genes, genes overlapping with the raw core and method-specific increments.

**Extend Data Fig. 11.**
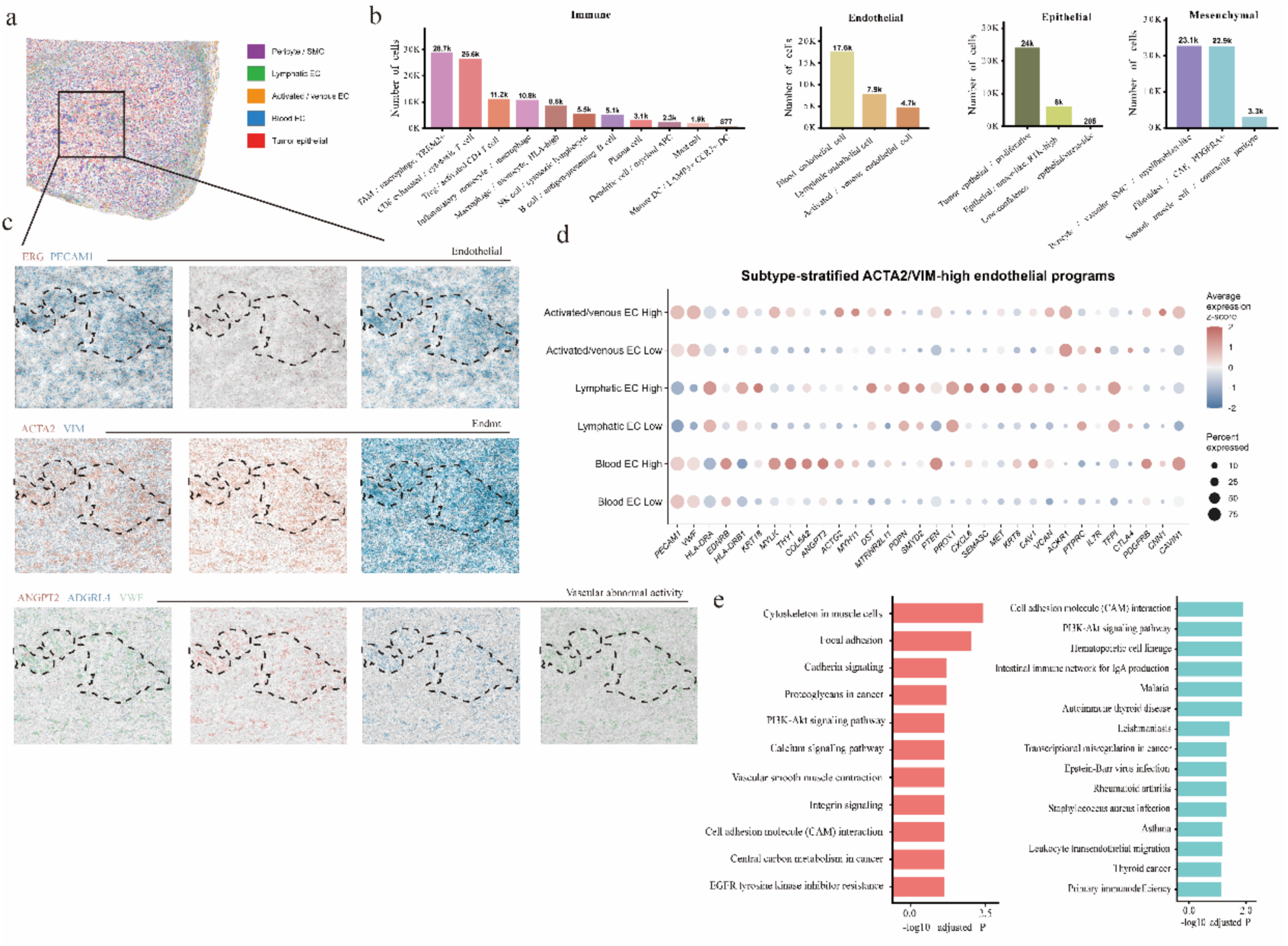
Spatial validation of EndMT-like endothelial states in renal cancer **a,** Spatial map of the renal tumour section, coloured by major epithelial, endothelial and mesenchymal compartments. **b,** Cell numbers for immune, endothelial, epithelial and mesenchymal subtypes identified in the spatial dataset. **c,** Spatial expression of endothelial markers ERG and PECAM1, EndMT-associated markers ACTA2 and VIM, and vascular-abnormality markers ANGPT2, ADGRL4 and VWF within the enlarged region. Dashed outlines mark representative vascular domains. **d,** Subtype-stratified endothelial programmes in ACTA2/VIM-high and ACTA2/VIM-low cells from activated or venous, lymphatic and blood endothelial populations. Dot size denotes the percentage of cells expressing each gene and colour denotes average scaled expression. **e,** Pathway enrichment in ACTA2/VIM-high endothelial populations. Left, programmes associated with cytoskeletal remodelling, focal adhesion, cadherin, PI3K– Akt, smooth-muscle contraction and integrin signalling. Right, immune- and adhesion-related programmes. Bars show −log10-adjusted P values.

